# Fuel buildup shapes post-fire fuel decomposition through soil heating effects on plants, fungi, and soil chemistry

**DOI:** 10.1101/2024.05.08.592975

**Authors:** Jacob R. Hopkins, Tatiana A. Semenova-Nelsen, Jean M. Huffman, Neil J. Jones, Kevin M. Robertson, William J. Platt, Benjamin A. Sikes

**Affiliations:** The Ohio State University; Evolution, Ecology & Organismal Biology; 318 W 12^th^ Ave Aronoff Laboratory, Columbus, OH 43210, USA; University of Kansas; Kansas Biological Survey, 2101 Constant Avenue, Takeru Higuchi Hall, Lawrence, KS 66047, USA; Tall Timbers Research Station; 13093 Henry Beadel Rd., Tallahassee, FL, USA; Louisiana State University; Department of Biological Sciences; 202 Life Science Bldg., Baton Rouge, LA 70803, USA; University of Kansas; Ecology & Evolutionary Biology; 1200 Sunnyside Avenue Haworth Hall, Lawrence, KS 66045, USA

**Keywords:** fire, decomposition, feedbacks, fuel accumulation, fungi, soil heating

## Abstract

Forty percent of terrestrial ecosystems require recurrent fires engineered by feedbacks between fire and plant fuels. Fuel loads control fire intensity which alters soil nutrients and shapes soil microbial and plant community responses to fire. Changes to post-fire plant fuel production are well known to feed back to future fires, but post-fire decomposition of new fuels is poorly understood. Our study sought to quantify how pre-fire fuel loading impacted post-fire fuel decomposition through soil abiotic properties, plant and soil fungal communities. In a longleaf pine savanna, both near and away from overstory pines, we manipulated pre-fire plot fuel loads to modify soil heating. We then assessed how fuel load and soil heating influenced post-fire plant fuel decomposition through changes to soil chemistry, vegetation, and fungi. Larger fuel loads, particularly beneath pines, increased soil heating and reduced decomposition of newly deposited fuels during the eight months following fire. Fire intensity effects on soil nutrients had the most consistent effects on decomposition with plant and fungal communities playing secondary roles. This demonstrates how fuel load and soil heating influence post-fire decomposition through fire-driven changes to soil abiotic properties, plant communities, and soil fungi. Further, since fire effects on decomposition and fire-fuel feedbacks were temporally dynamic this illustrates the importance of considering fire-fuel feedbacks across time. Understanding the importance of these feedbacks among ecosystems can help increase our predictive ability to manage fuels and the effects of repeated fires.

## 1. Introduction

Fire-dependent ecosystems are maintained by feedbacks between fire and fuels. Recurrent fires maintain >40% of Earth’s terrestrial ecosystems (Archibald et al., 2013; Bowman et al., 2009). When fire is excluded, biodiversity declines (Harms et al., 2017; He et al., 2019; Neary and Leonard, 2020) and systems shift to other ecosystem states (Beckage et al., 2011, 2009; Beckett and Bond, 2019; Tiribelli et al., 2018). Fire effects in these “pyrophilic” ecosystems are largely determined by fire history, frequency, and intensity (Alcañiz et al., 2018; Certini, 2005; He et al., 2019; Pressler et al., 2019), with fuel accumulation directly relating to fire’s impact on vegetation and soils (i.e., fire severity). Vegetation produces fuels for fires, therefore plant responses to fire create a feedback to the buildup of new fuels for future fires (i.e., fire-fuel feedbacks; Jensen *et al*., 2001; Hart *et al*., 2005; Beckage *et al*., 2009; Platt *et al*., 2015). Fires also alter fuel loads through changes to decomposition in both rarely burned (Holden and Treseder, 2013; Pellegrini et al., 2020) and pyrophilic ecosystems (Brennan et al., 2009; Ficken and Wright, 2017; Hopkins et al., 2020; Throop et al., 2017). While significant work has explored relationships between plant fuel production, increasing fuel loads, and fire intensity, little work has quantified the pathways for feedbacks between pre-fire fuels and post-fire decomposition of new fuels. Reintroducing prescribed fires to restore and maintain threatened pyrophilic ecosystems demands a clear understanding of these fire-fuel feedbacks to tailor fire regimes and provide effective management (Davies et al., 2019; Lake et al., 2017; Scott et al., 2012; Varner et al., 2005).

By influencing fire intensity, fuel loads can result in specific ecological outcomes that may determine fire-fuel feedbacks. Ecosystem differences in vegetation flammability, fuel accumulation, and weather conditions all give rise to variation in fuel loads and therefore fire characteristics (Archibald et al., 2013; Bowman et al., 2009). Woody fuels, for example, generally make savanna fires more severe than those in grasslands (Ford, 2010; Platt et al., 2016a). Hotspots of fire severity may be localized within an ecosystem too, for example near bunchgrasses or where oak leaves or pine needles accumulate below canopy trees (Beckage et al., 2011, 2009). This variation is a core component of pyrodiversity (Hu et al. 2019), the heterogenous landscape effect of fire regimes that helps foster biological diversity. Where more fuels accumulate, fuel producers (i.e., plants) *and* decomposers will experience more intense fires while areas with few fuels may act as fire refugia for more sensitive species (Beckett and Bond, 2019; Meddens et al., 2018). Fire severity variation also impacts abiotic soil properties and these changes indirectly shape the post-fire responses of vegetation and litter decomposers (Day et al., 2020; Gibb et al., 2022; Neary and Leonard, 2020; Resco de Dios, 2020). Combined, these abiotic and biotic pathways affect fuel buildup, and thus fire intensity, which may produce predictable effects on post-fire fuel loads.

The role of soils in fires is complex but little experimental work has quantified how their responses to fire severity can feedback to the fuel cycle. Fine fuels and organic matter combustion generally increase soil pH and release nutrients in the short term (Butler et al., 2018; Certini, 2005), particularly in grasslands and savannas with low intensity fires (Alcañiz et al., 2018; Ford, 2010; Neary and Leonard, 2020). Longer, hotter fires caused by fallen trees can drive a net loss of nitrogen and carbon through volatilization and combustion (Johnson and Curtis, 2001; Raison, 1979). Edaphic shifts in both cases can influence post-fire plant composition and fuel production (Gagnon et al., 2015; Harms et al., 2017; Hopkins et al., 2023), as well as the microorganisms that underlie them. Soil biota, including bacteria and fungi also show differential responses to fire intensity, with larger declines from high severity fires where heat penetrates deeper in the soil (Caiafa et al., 2023; Peay et al., 2009; Taudière et al., 2017). Many fungi appear particularly sensitive to fire, more so than many bacteria (Bárcenas-Moreno and Bååth, 2009; Dooley and Treseder, 2012; Hamman et al., 2007; Pulido-Chavez et al., 2023), with sensitive groups including fungal mutualists, on which plant productivity depends, and saprotrophic fungi, whose fire responses regulate fuel longevity (Fox et al., 2022; SemenovaLJNelsen et al., 2019). Local variation in fuel loads and thus soil heating may affect the interactions between soil abiotic and biotic components which ultimately shape fire-fuel feedbacks. Moreover, because fire intensity sets the clock for post-fire effects, the timing of feedbacks among these different system components may be particularly important to quantify but remains largely unexplored.

Here we tested how fuel load and associated soil heating influence the decomposition of post-fire plant fuels through plant, soil, and soil fungal changes in a longleaf pine ecosystem. As in most savannas, pyrophilic plants form a continuous layer of surface vegetation and fire-resistant trees (here longleaf pines) form an incomplete canopy (Peet et al., 2018). The spatial heterogeneity of the pine canopy creates variation in fire severity through differences in pine needle and grass fuel loads (Ellair and Platt, 2013; Platt et al., 2016b; Sánchez-López et al., 2023). Previous work in this system has linked variation in fire severity and frequency to changes in fungal (SemenovaLJNelsen et al., 2019) and bacterial communities (Dao et al. 2022), as well as reduced decomposition (Hopkins et al., 2020). In this experiment, we manipulated fine fuels in plots near and away from canopy pines to test how fuel loads and heat release influenced soil conditions, plant communities, fungal communities, and the microbial decomposition of post-fire plant fuels. We hypothesized that larger fuel loads and resulting increases in soil heating during fire would decrease microbial decomposition of post-fire fuels (Fig. 1), because of greater fire effects on vegetation and soils. Our findings suggest that fire effects on microbial decomposition are mediated by interactions between vegetation and belowground systems.

**Figure 1:**
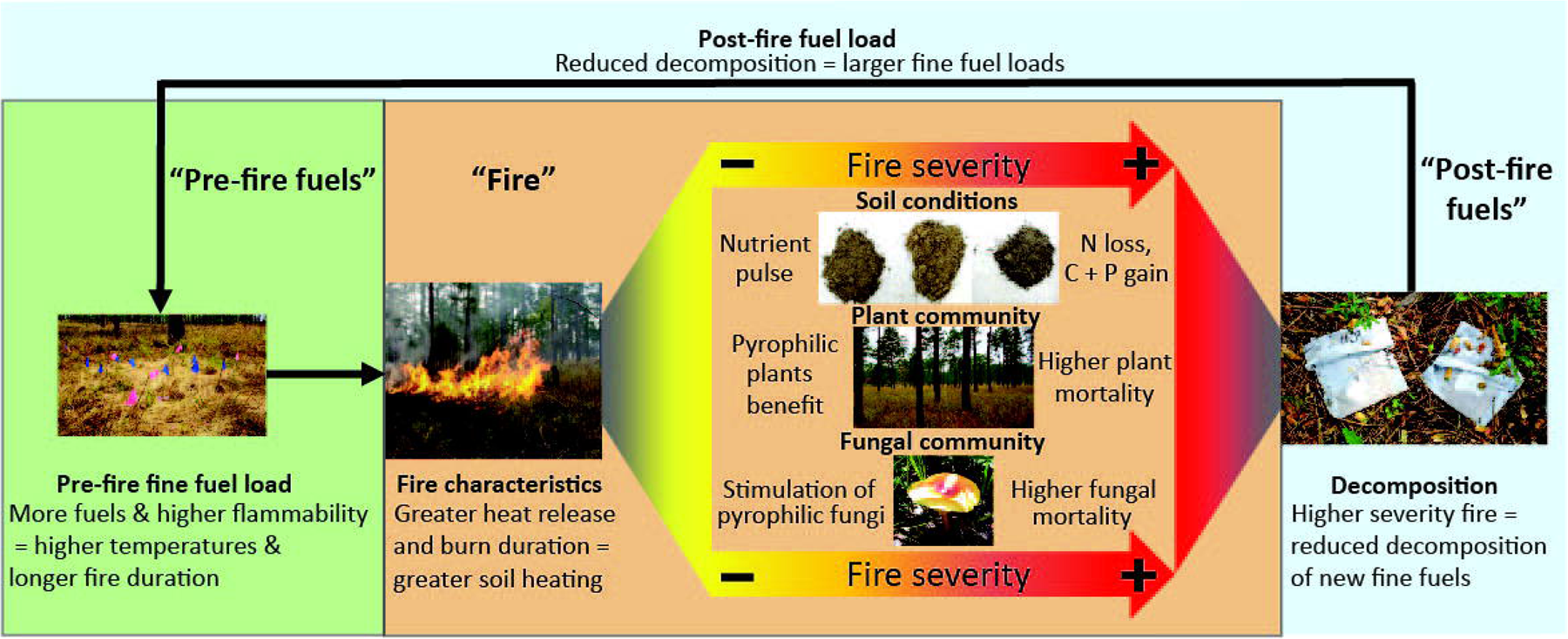
Hypothesized pathways through which fine fuel load influences the microbial decomposition of post-fire fine fuels. Pre-fire fine fuel mass and flammability determine fire characteristics associated with greater heat release, combustion, and soil heating. As soil heating increases, fire becomes more severe and its influence on soil conditions, vegetation, and soil fungi changes. Depending on soil heating effects on these components (i.e., severity), post-fire microbial decomposition of new fine fuels may be affected, and alter post-fire fine fuel loads.

## 2. Materials and Methods

### 2.1. Field Site

This study was conducted on the Wade Tract in Thomas County, GA, USA (30° 45’ N; 84° 00’ W; Platt *et al.,* 1988, *Peet et al.,* 2018; Figure S1). This site has a 10-month growing season with an average precipitation of 1350 mm. Soils are Typic and Arenic Kandiudult Ultisols characterized by sand or loamy sand A and E horizons with sandy clay loam Bt sub horizons (USDA NRCS, 2023). Two prescribed fire units containing an old-growth conservation easement were used. All fire units have historic fire return intervals of 1-3 years, and have been managed for the past century with annual/biennial prescribed fires mostly between January and June (Rother et al., 2020).

### 2.2. 2017 Field Plots

This study took place within the year following prescribed fires conducted in 2017. The two fire management units were burned on March 23^rd^ (East) and April 12^th^ (West) using drip torches and flanking fires ignited in late morning, with relative humidities of 37-83%. Flame lengths generally ranged from 0.5 to 2.5 m, and average fuel consumption by fire was estimated at 55% in an adjacent study (Reid et al., 2012).

We used GPS maps indicating locations of *Pinus palustris* canopy trees (developed by Robertson *et al*., 2019) to choose eight experimental patches greater than 5 m^2^, four of which were located within 10 m of overstory longleaf pines (near), and the other four were at least 10 m away from the nearest overstory pine (away). Within each patch, twelve 4 m^2^ plots were established and received one of five fire severity treatments (described below). This produced 96 total plots, 48 near and 48 away from overstory pines.

### 2.3. Fuel load Treatments

Five fuel load treatments were created by manipulating pyrogenic pine needle fuels that influence fuel consumption, heat release, and soil heating (Platt et al. 2016). Fuel treatments were randomly applied to plots in each patch. The treatments were meant to produce fires with low to high total heat release like those that occur naturally on the landscape: removal of pine needles (removal), no manipulation of pine needles (reference), swapping proportions of pine needle and grass fuels between near and away plots (switch), 1000 g/m^2^ addition of pine needles (1X), and 2000 g/m^2^ addition of pine needles (2X). Removal treatments produced fine fuel beds limited to surface layer vegetation. Reference treatments represented natural pine needle accumulation. Switch treatments allowed us to assess whether fuel characteristics, including flammability, in addition to fuel mass governed fire characteristics. 1X and 2X treatments mimicked fuel loads under high pine densities or fallen pine crowns. Because of existing fuel heterogeneity in the field, the specific fuel quantities for removal and addition were tailored to existing fuels in each patch location as detailed in Figure S2. After the 2017 fires, we mapped the spatial extent of fires with GPS then located 10 adjacent patches that did not burn (no burn), for use as comparisons to burned plots.

### 2.4. Fuel and Fire Characteristic Analyses

Fuel loads were sampled before and after 2017 prescribed fires. One week prior to 2017 fires, paired 30 x 30 cm subplots were established in each plot. Fine fuels (or residuals) were collected, dried, and weighed in one of the subplots before and the other after fire. Fuel consumption was estimated as the difference between pre-fire and post-fire fuels.

Characteristics of the 2017 fires were measured using established procedures (Ellair and Platt, 2013; Gagnon et al., 2015). Two, 0.81 mm thick thermocouples (XCIB-K-2-3-10; Omega Engineering, Inc., Stamford, CT) were placed in the center of each plot: 1cm above the ground surface (not contacting litter or soil) to estimate surface heating and combustion residence time and the other 1 cm below the soil surface to measure soil heating. Thermocouples were attached to U12-014 data loggers (Omega Engineering) which recorded temperatures every second from the time of activation until collected several hours after passage of flaming fronts of the prescribed fires. Maximum litter and soil temperature were calculated as the highest temperature spikes recorded by the litter and soil thermocouples during fire. Duration of heating was calculated as the time (seconds) that the litter temperature remained >60°C, a widely used threshold for lethal effects on plant and microbial tissue (Fox et al., 2022; Platt et al., 2016a).

### 2.5. Quantifying Microbial Decomposition

To measure microbial decomposition, 5 kg of recently deposited, intact dead plant biomass was collected outside the 4 m^2^ unburned plots in March 2017. This new litter included pine needles, grass culms, forbs, and oak leaves produced that year; partially decomposed litter on the ground surface was not included. Collected litter was kept separate from unburned patches near and away from pines. All collected litter was shipped to the University of Kansas where it was stored at 4°C for less than 1 week until processed. The plant litter was dried at 65°C for 72 hours, ground using a Model 4 Wiley Mill (Thomas Scientific, Swedesboro, NJ) with a 6 mm opening, and then sterilized via gamma irradiation to ∼32 kGy at the Penn State Radiation Science & Engineering Center. In a biological safety cabinet, sterilized plant litter was placed in 15 x 15 cm, 30 µm nylon mesh bags (Robertson & Paul 2000), a mesh size which isolates microbial decomposition (Bradford et al., 2002). Each decomposition bag was filled with approximately 4 g of plant litter, collected from either near or away from pines. Initial mass for each bag was recorded before and after litter was added; then bags were heat sealed and stored in sterile plastic bags until deployment in plots.

Bags were placed in the plots in May 2017, approximately 1-2 months after fire. Four decomposition bags, each with litter that corresponded to local pine overstory conditions (e.g., near litter for near pine plots), were randomly selected and placed on the soil surface in each plot. Bags were placed among the bases of vegetation in plots so the flat surface of the bag contacted litter and soil, then anchored along margins with 5 cm sod-staples to ensure that they remained in place. Bags were deployed after the 2017 prescribed fires and collected 2, 4, 6, and 8 months post-deployment. Any surface soil or litter on the bags was carefully removed, then bags were placed in sterile plastic bags and shipped overnight to the University of Kansas (Lawrence, KS) for analysis. For each bag, litter content was carefully removed, dried at 65°C for 72 hours, and then weighed to determine mass loss. Decomposition rates (k) were then calculated for each bag as:

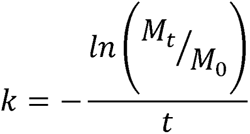

where M_0_ is the starting mass, M_t_ is the mass at the time of collection, and t is the number of days which the bag was deployed in the field. Use of k-rates allows for the downstream use of linear methods when analyzing decomposition (Karberg et al., 2008; Olson, 1963).

### 2.6. Soil Analyses

To assess fuel manipulation and fire effects on soil chemistry, we collected soil from the upper 5 cm of each plot 2 months after 2017 fires. Soil was kept cool in the field and shipped to the University of Kansas and stored at 4° C until analysis (2 weeks). Soils were then sent to the Kansas State University Soil Testing Lab, dried and ground prior to analysis, and analyzed for total phosphorous, total carbon, total nitrogen, ammonium, nitrate, C:N ratios, and pH. Soil P was analyzed using the Mehlich-3 method on a Lachat Quickchem 8000 (Lachat Instruments, Loveland, CO; Mehlich, 1984). Total C and N were quantified on a LECO TruSpec CN Carbon/Nitrogen combustion analyzer (LECO Corp., St. Joseph, MI. NH_4_ ^+^ and NO_3_ ^-^ were extracted with 2 M KCl per 2 g of soil, and a cadmium reduction for nitrate and colorimetric procedures were used, followed by flow analysis (Brown 1998). Soil pH was measured by averaging the pH of three separate soil samples per plot 1:1 with pH 7 H_2_O. C:N ratio was estimated by dividing total C by total N.

### 2.7. Plant community Sampling

To assess fire effects on vegetation, we quantified plant community composition in July and August following 2017 prescribed fires. A 1 m^2^ subplot was established in each experimental plot. All plant species within these subplots were recorded (presence/absence) and species richness was quantified.

### 2.8. Sampling of Fungal Communities

Litter and the uppermost soil layer inside each plot were sampled two weeks before and two months after 2017 prescribed fires. Samples were carefully collected from two randomly selected 9 x 9 cm^2^ areas in each plot. First, litter was collected manually from the soil surface, ensuring that no recently fallen material (post-fire) was collected. In burned plots, litter was often charred, but was not entirely consumed in the most recent fire; in unburned plots, litter had been present for some time, typically since the previous fall. After litter was collected, surficial soils to a depth of 1.5 cm were collected. The litter and soil collected from each plot were pooled in separate plastic bags. To avoid cross-contamination, all sampling equipment was sterilized between plots; first with 10% bleach and then with 90% isopropyl alcohol. Soil and litter samples were kept cool in the field, frozen at -20°C within 4 h, and then shipped overnight to the University of Kansas, where they were stored at -80°C. Prior to DNA extraction, soil samples were homogenized and subsampled twice (e.g., 250ml to 25ml to 2.5ml) to help ensure community data were not biased by sampling effects. Litter was ground under liquid nitrogen prior to subsampling.

### 2.9. DNA Extraction and PCR

To analyze fungal community composition, DNA was extracted from 0.25 g soil and litter subsamples using NucleoSpin® Soil Kits (Machery-Nagel, Düren, Germany) following the manufacturer’s protocols. A single-step PCR was used to amplify the ITS2 region with the fITS7 (forward) and ITS4 (reverse) primers (Ihrmark et al. 2012, White et al. 1990) and a PCR mix of 1.0 µL of DNA, 8 µL of 5 x Q5® buffer (New England Biosystems, Ipswich, MA), 0.8 μL of dNTPS (10 mM), 2 μL of each primer (10 mM), 0.4 μL of Q5® High-Fidelity DNA polymerase (New England Biosystems), 8 μL of enhancer (New England Biosystems), and 17.8 µL of dd H2O for a reaction volume of 40 µL. The PCR scheme had an initial denaturation step at 98° C for 30 sec, then 25 cycles of 98° C for 10 sec, 57° C for 30 sec, and 72° C for 30 sec, a final extension step at 72° C for 2 min, and a hold at 4° C. PCR products were checked on agarose gels to ensure amplification, then cleaned using Agencourt AMPure XP magnetic beads (Beckman Coulter, Indianapolis, IN).

### 2.10. Library preparation and Sequencing

We used the Illumina MiSeq Nextera protocol to sequence litter and soil fungal community samples. With a second PCR reaction, sequence barcodes (Nextera indices, Illumina, San Diego, CA) were added to the samples. The second PCR followed the first, except 5 µL of PCR amplicon was used, and PCR cycles were set to eight. Barcode amplicons were purified with Agencourt beads and DNA concentration was measured using a Qubit 2.0 (LifeTechnologies, Carlsbad, CA). Samples were then pooled in equimolar concentrations and sequenced using an Illumina MiSeq (Illumina, San Diego, CA) with V3 chemistry at the Kansas State Integrated Genomics Center. Sequence data is deposited in the GenBank Sequence Read Archive (SRA) PRJN#####.

### 2.11. Bioinformatics

Sequence data were analyzed using Qiime2 version 2019.1 using methods described in Bolyen et al. 2019. Quality and barcode filtering resulted in 5M reads. Unique barcodes were trimmed from paired reads using cutadapt (Martin, 2011), then denoised, merged and chimera checked using the dada2 tool (Callahan et al., 2016) implementation within Qiime2. The UNITE fungal ITS reference database v8 “dynamic” (Abarenkov *et al*., 2010; accessed April 2020) was used to train a Naïve Bayes classifier, which defined amplicon sequence variants (ASVs) and assigned taxonomic identities. ASVs with less than five reads were removed to conservatively separate data from noise.

### 2.12. Data Analyses

All analyses were conducted in R version 4.1.3 (R Core Team 2022). We tested how fire characteristics (temperature at the surface, in soil, and duration above 60°C) and fine fuel combustion varied among fuel loading treatments using type III multivariate analysis of variance (MANOVA) with the base manova() function and the joint_tests() function in the emmeans package (Lenth, 2018). The model included fuel loading treatment, pine proximity, and their interaction as fixed effects, as well as a management unit term to control for unmeasured differences between the two prescribed burns. Estimated marginal means were extracted using the emmeans() function, and treatment differences assessed using the contrast() function. Fire characteristic variables were natural log transformed using the log() function to help model residuals meet assumptions.

Differences in soil chemistry among experimental treatments were assessed similarly to fire characteristics and fine fuel combustion. Total inorganic phosphorous and nitrate values were log transformed to meet model assumptions.

Plant community composition differences among fuel loading and pine proximity treatments were tested with principal coordinates analysis (PCoA) and permutational multivariate analysis of variance (PERMANOVA) in the vegan package (Oksanen et al., 2013). The plant community matrix was transformed using the Bray-Curtis method using the vegdist() and princomp() functions. After ordination, a PERMANOVA containing fuel load, pine proximity, and their interaction, as well as fire management unit and patch (nested in fire unit) terms was used to assess treatment effects on plant community composition. Differences in plant communities among treatments were assessed with the pairwise.perm.manova() function in the RVAideMemoire package (Hervé, 2021). Plant community richness was quantified using the specnumber() function. Differences in plant community richness were tested using type III linear mixed effect models (LMERs) with the lmer() function in the lmer package (Bates et al., 2015). The richness LMER model was identical to the MANOVA models for fire above, however, management unit and patch (nested within unit) were included as random effects.

Fuel loading and pine proximity effects on microbial decomposition rate (k) were tested using MANOVAs and contrasts identical to those for fire characteristics and soil factors. These analyses include assessment of overall k-rates and variation at individual time points.

Differences in soil and litter fungal community composition were assessed similarly to plant communities, using PCoAs and PERMANOVAs. First, excess zeroes in the fungal ASV matrix were replaced using the cmultRepl() function with the Bayesian Laplace method available in the zCompositions package (Palarea-Albaladejo and Martín-Fernández, 2015), then the fungal ASV matrix was center-log ratio transformed and Aitchison’s distance used to create a dissimilarity matrix within the vegan package (Quinn et al., 2019). After ordination, we ran a PERMANOVA testing effects of fuel loading treatment, pine proximity, sampling time (pre vs. post), community type (litter vs. soil), their interactions, and management unit on fungal community composition. Contrasts were assessed similarly to plant communities. Additionally, post-fire fungal diversity (inverse Simpson metric) and community heterogeneity (i.e., inter-plot community variation) were quantified using the diversity() and betadisper() functions in the vegan package and tested using a LMER model identical to those above and ANOVA model with fuel loading treatment as a fixed effect, respectively. Finally, the ALDEx2 package (Fernandes et al., 2013) was used to identify fungal indicator species with differential abundance between pre- and post-fire communities for each fuel loading treatment. Indicator species p-values were adjusted using the Benjamini-Hochberg method.

We explored the mechanisms mediating fuel loading effects on microbial decomposition using structural equation modeling (SEM) with the lavaan package (Rosseel, 2012). We hypothesized a model that linked fire effects on plant communities, soil chemistry, and fungal communities, to the microbial decomposition of fine fuels (Figure 2; Tables 1, S3). This model first created a latent variable representing soil heating effects using the measured fire characteristics (Figure 2, path a), then explored its’ direct and indirect effects on plant community composition, soil chemistry, and fungal community composition (Figure 2, path b). We further hypothesized that fire would influence microbial decomposition through changes to soil factors, plants and fungal communities (Figure 2, path c). Model fitting was guided by a goodness of fit and parsimony approach, where poorly supported paths were iteratively removed (e.g., p > 0.75, p > 0.5, and p > 0.25; Table S4). Pre-fire and no burn samples were omitted from SEMs and continuous variables were scaled prior to analysis. The final model is detailed in appendix section 3.

**Table 1:** SEM variable description table. Contains the means, standard deviations, and type information for each variable used in SEM analyses. Variables are the same as those used in univariate analyses. Continuous variables were scaled prior to SEM analysis.

| model term | variable type | mean | s.d. | description |
| --- | --- | --- | --- | --- |
| max. surf. temp. | °C | 356.14 | 228.3281 | Max. temp. change (°C) in litter layer |
| surf. dur. >60C | sec | 813.579 | 2283.807 | Time (seconds) fires remained over 60°C |
| max. soil temp. | °C | 48.24 | 56.36419 | Max. temp. change (°C) in soil layer |
| fuel manipulation | factor | 0 = no burn,<br>1 = removal, 2<br>= reference, 3<br>= switch, 4 =<br>add 1x, 5 =<br>add 2x | - | Fuel manipulation treatment in plot |
| pine proximity | factor | 0 = away, 1<br>= near | - | Near = within 10m of nearest pine, Away<br>= 10 m away of nearest pine |
| plant PCoA axis 1 | cont. | 0.11 | 0.669108 | Post-fire plant community composition |
| plant PCoA axis 2 | cont. | -0.04 | 0.360356 | Post-fire plant community composition |
| plant species richness | cont. | 23.671 | 4.636893 | Post-fire plant species richness |
| inorg. phosp. | ppm | 34.65 | 32.73722 | Determined by Mehlich III method |
| C:N ratio | ratio | 27.868 | 4.596712 | Ratio of total soil carbon to nitrogen |
| soil pH | cont. | 6.321 | 0.280177 | Soil pH in each plot |
| fine fuel combustion | percent | 74.501 | 23.96209 | Fuel consumed during prescribed fires |
| litter/soil fungi PCA1 | cont. | 0 | 163.8966 | Post-fire litter/soil fungal comm. |
| litter/soil fungi PCA2 | cont. | 0 | 26.58693 | Post-fire litter/soil fungal comm. |
| litter/soil fungal<br>diversity | cont. | 69.565 | 41.2983 | Post-fire litter and soil fungal inverse<br>Simpson metrics |
| month two decomp. | cont. | 0.002267 | 0.000759 | Decomp. rate (k) two months post-fire |
| month four decomp. | cont. | 0.002201 | 0.000776 | Decomp. rate (k) four months post-fire |
| month six decomp. | cont. | 0.001956 | 0.000792 | Decomp. rate (k) six months post-fire |
| month eight decomp. | cont. | 0.001628 | 0.000617 | Decomp. rate (k) eight months post-fire |

**Figure 2:**
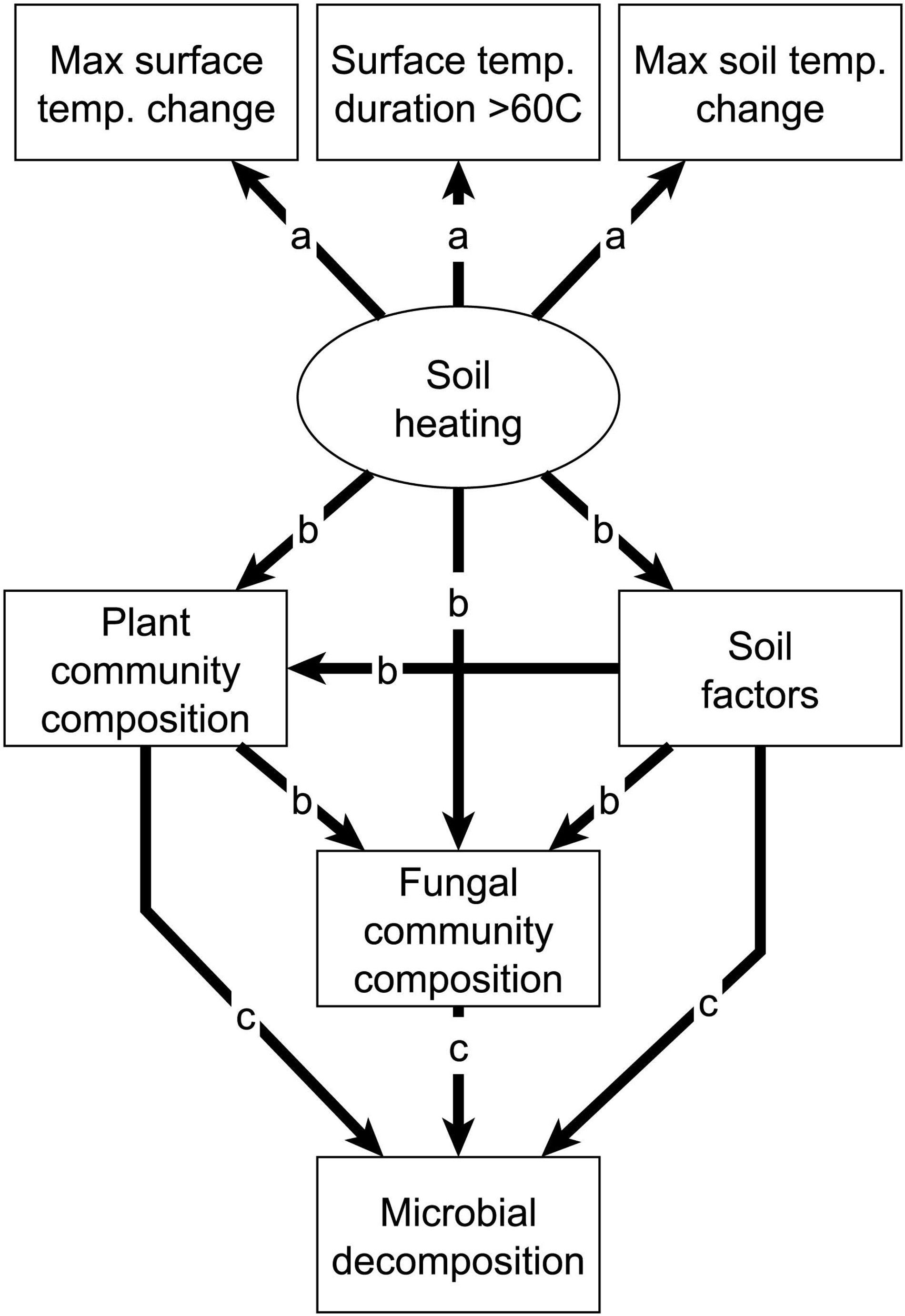
Hypothesized pathways for soil heating effects on microbial decomposition: a) fire characteristics like maximum temperature change and duration >60° C reflect soil heating, b) soil heating alters soil factors (C, N, P, pH), plant community composition, and fungal community composition directly and indirectly, which in in turn c) shapes microbial decomposition of post-fire fine fuels.

## 3. Results

### 3.1. Fuel manipulations altered combustion and soil heating as intended

The fuel load treatments generated overall differences in soil heating and fine fuel combustion (F_4,66_=15.2, p<0.0001; Fig. S6, Tables 2 & S5), with the 1X and 2X addition plots showing the greatest heating effects. Individually, fire characteristics (temperature and heating duration) and fine fuel combustion also varied in response to fuel manipulation treatments (F_12,66_=5.76, p<0.0001). As more fuels were added, maximum surface temperatures (Figure S6a) and durations >60° C increased (Figure S6b), although these increases were primarily in contrast to the fuel removals. Maximum soil temperature changes (Figure S6c) and fine fuel combustion (Fig. S6d) also increased predictably with more fuels, with the largest values in the 1X and 2X treatments. Increasing fuels in near pines plots generated larger temperature changes but did not in away from pines plots. In summary, fuel manipulations predictably influenced soil heating.

**Table 2:** MANOVA results for fuel manipulation and pine proximity effects on 2017 prescribed fire and fine fuel combustion characteristics.

| model term | d.f. 1 | d.f. 2 | F-test | p-value |
| --- | --- | --- | --- | --- |
| *fuel manip. treatment* | 4 | 66 | 15.239 | <0.0001*** |
| *pine proximity* | 1 | 66 | 1.779 | 0.1868 |
| *fire unit* | 1 | 66 | 8.419 | 0.005** |
| *repeated measure* | 3 | 66 | 511.292 | <0.0001*** |
| *fuel x pine* | 4 | 66 | 1.473 | 0.2203 |
| *fuel x rep. meas.* | 12 | 66 | 5.76 | <0.0001*** |
| *pine x rep. meas.* | 3 | 66 | 1.808 | 0.1542 |
| *unit x rep. meas.* | 3 | 66 | 5.129 | 0.003** |
| *fuel x pine x rep. meas.* | 12 | 66 | 0.891 | 0.5597 |

. = $p < 0.1$ ; \* = $p < 0.05$ ; \*\* = $p < 0.0001$

### 3.2. Soil chemistry varied in response to fuel manipulations

Fuel manipulation effects varied among individual soil properties (F_30,66_=3.01, p=0.0001; Table 3, S8-S9). Larger fuel loads (and thus hotter fires) increased inorganic phosphorus (Fig. S7a), ammonium (Fig. S7b), and soil pH (Fig. S7c), although pH changes only occurred as fuel loading increased near pines (F_5,63.6_=2.92, p=0.02). Total nitrogen total carbon, C:N ratios, and nitrate (Fig. S7d-g) did not respond to fuel manipulation treatments or proximity to overstory pines.

**Table 3:** MANOVA results for fuel manipulation treatment and pine proximity effects on soil chemistry.

| model term | d.f. 1 | d.f. 2 | F-test | p-value |
| --- | --- | --- | --- | --- |
| *fuel manip. treatment* | 5 | 66 | 1.128 | 0.3543 |
| *pine proximity* | 1 | 66 | 2.039 | 0.158 |
| *fire unit* | 1 | 66 | 2.671 | 0.107 |
| *repeated measure* | 6 | 66 | 13235.32 | <0.0001*** |
| *fuel x pine* | 5 | 66 | 0.717 | 0.6131 |
| *fuel x rep. meas.* | 30 | 66 | 3.011 | 0.0001*** |
| *pine x rep. meas.* | 6 | 66 | 1.651 | 0.1473 |
| *unit x rep. meas.* | 6 | 66 | 15.307 | <0.0001*** |
| *fuel x pine x rep. meas.* | 30 | 66 | 1.596 | 0.0582 |

\* $p < 0.05$ , \*\* $p < 0.01$ , \*\*\* $p < 0.0001$

### 3.3. Fuel manipulation altered plant community composition

Plant community composition independently varied with fuel manipulations (F_5,81_=1.61, p=0.004; R^2^=0.07; Fig. 3a, Table 4, S10) and proximity to overstory pines (F_1,81_=2.37, p=0.009; R^2^=0.02; Fig. 3b). Fire-related differences in plant composition were only between burned and unburned plots, however, not among burned treatments with different fuel loads. Plant species richness was also affected by fuel manipulations (F_5,37.1_=13.5, p<0.0001; Fig. 3c, Table S12, S13), with richness decreasing in higher fuel load treatments. Pine proximity did not influence plant species richness (F_1,16.1.1_=16.1, p=0.549).

**Figure 3:**
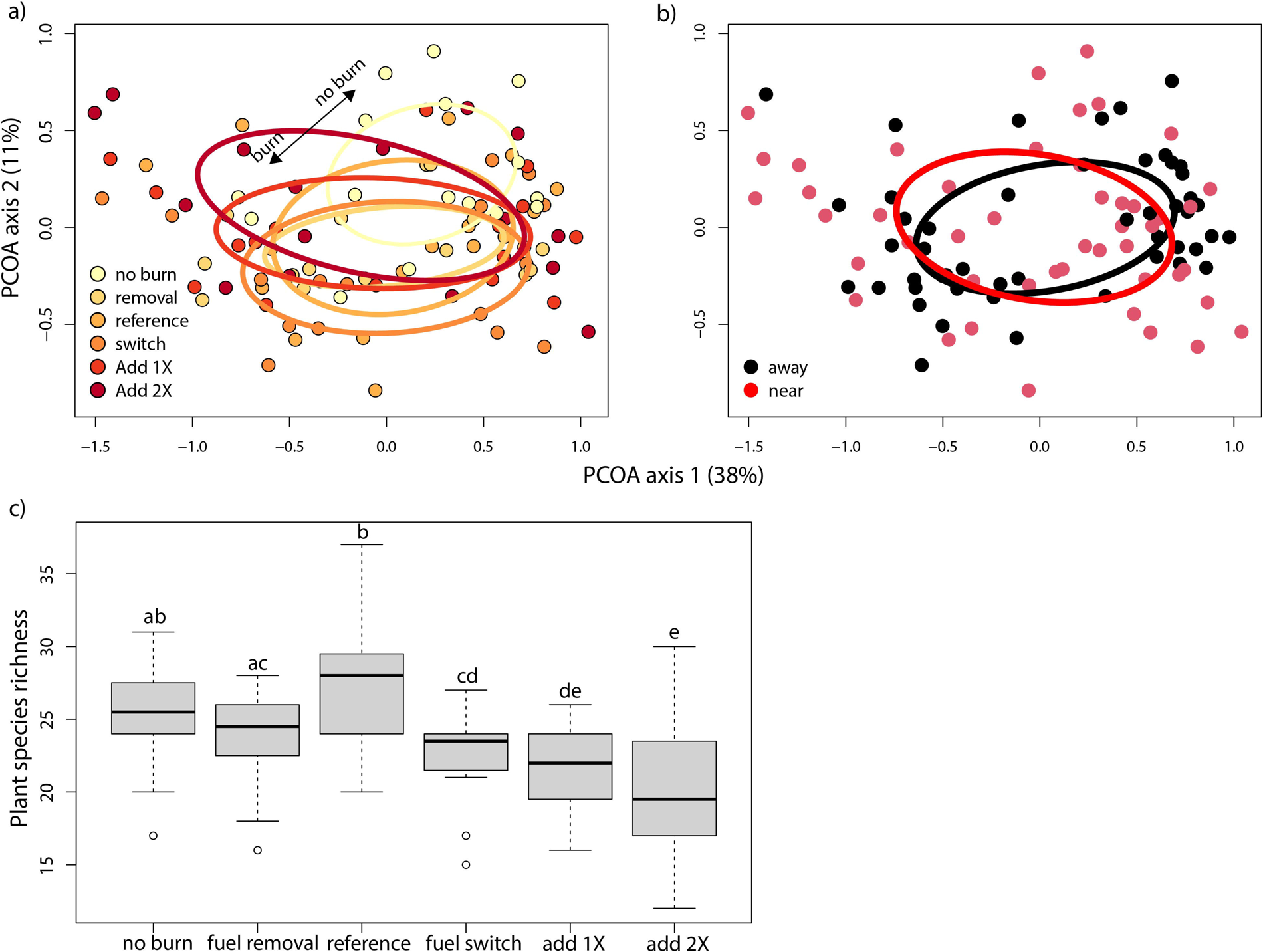
Fuel load treatment and pine proximity effects on plant communities. Ellipses in panel a and b represent 95% confidence intervals. Lower case letters in panel c represent differences between fuel load treatments. Following 2017 prescribed burns, a) plant communities in burned plots differed from those in nearby no burn plots, b) plant communities independently varied by proximity to overstory pines, and c) increasing pre-fire fuels decreased plant species richness.

**Table 4:** PERMANOVA results for fire severity treatment and pine proximity effects on plant community composition.

| Model term | d.f. | Sum Sqs. | Mean Sqs. | F-test | R<sup>2</sup> | p-value |
| --- | --- | --- | --- | --- | --- | --- |
| *fuel manip. treatment* | 5 | 1.1784 | 0.23569 | 1.6078 | 0.06963 | 0.0043 ** |
| *pine proximity* | 1 | 0.3477 | 0.34773 | 2.372 | 0.02055 | 0.008699 ** |
| *fire unit* | 1 | 1.6774 | 1.67743 | 11.4426 | 0.09911 | <0.0001*** |
| *fuel \* pines* | 5 | 0.6688 | 0.13377 | 0.9125 | 0.03952 | 0.664234 |
| *patch within fire unit* | 2 | 1.1778 | 0.5889 | 4.0172 | 0.06959 | <0.0001*** |
| *residuals* | 81 | 11.8742 | 0.14659 |  | 0.7016 |  |

\*p < 0.05, \*\*p < 0.01, \*\*\* p < 0.0001

### 3.4. Fuel manipulation altered fungal community composition

Fungal community composition varied among sampling times (pre vs. post), community types (litter vs. soil), fuel treatments, and pine proximities (F_4,174_=1.17, p=0.009; R^2^=0.017; Fig. 4, Table 5, S13-14). Prior to 2017 prescribed fires, fungal community composition differed between litter and soil communities, and based on proximity to overstory pines. Following prescribed fire, fungal communities in all fuel manipulation treatments (except for no burn plots) differed from pre-fire communities and displayed distinct variation among fuel treatments. There was significant spatial variation as location explained the single most variation among fungal communities (R^2^=0.106, Table 5). Fire effects on fungal communities were similar between near and away from pine treatments.

**Figure 4:**
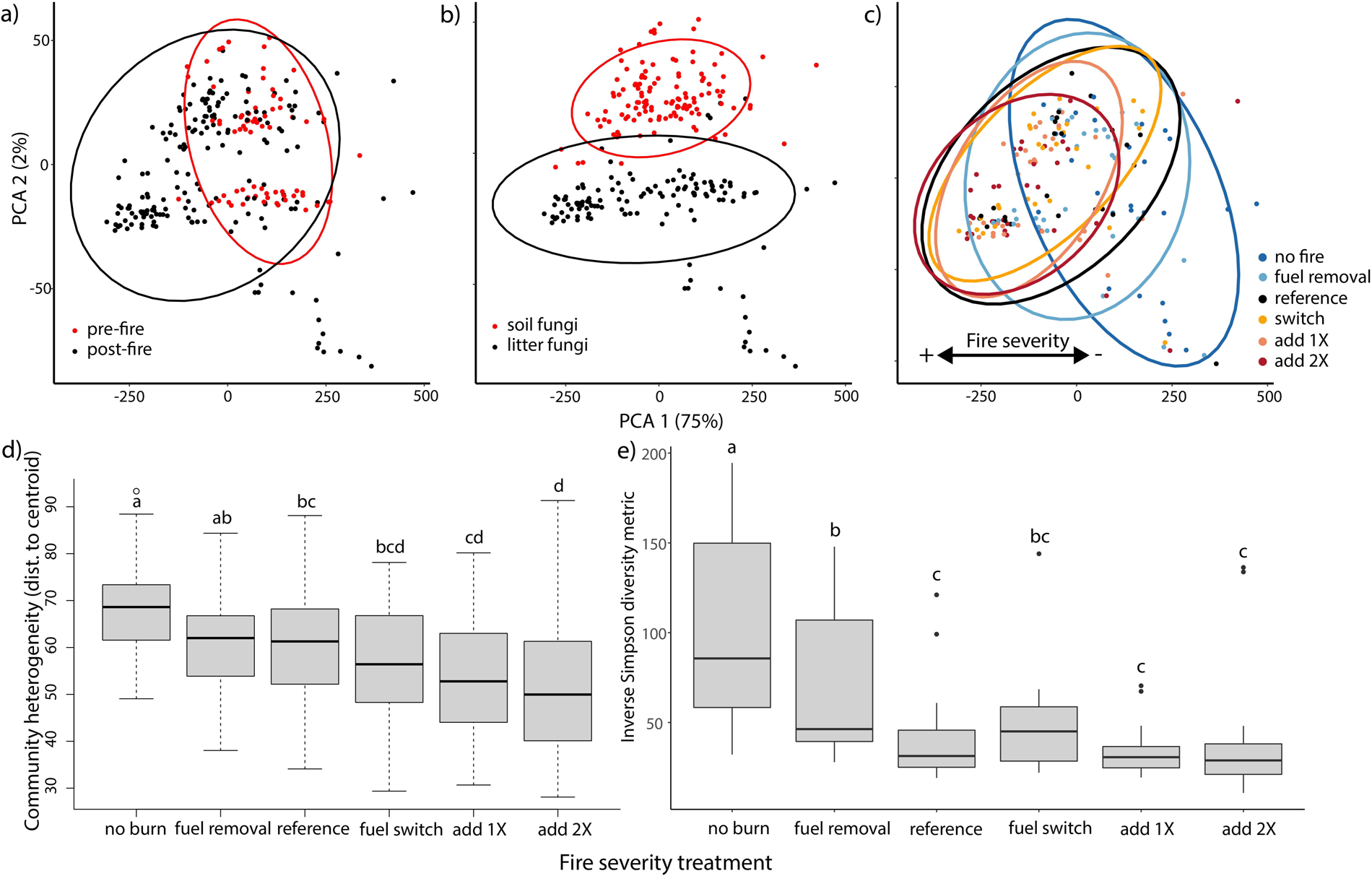
Fuel load and pine proximity effects on fungal community composition. Ellipses in panels a-c represent 95% confidence intervals. Lower case levels in panels d and e represent differences between fire load treatments. Fungal communities varied a) pre and post fire, b) between litter and soil samples, as well as c) fire fuel load treatments. Soil heating driven shifts in fungal community composition were associated with decreases in d) community heterogeneity and e) diversity.

**Table 5:** PERMANOVA model for fuel manipulation, pine proximity, community type (litter vs. soil), and sampling time (pre vs. post fire) effects on fungal community composition.

| Model term | d.f. | Sum Sqrs. | Mean Sqrs. | F-test | R2 | p-value |
| --- | --- | --- | --- | --- | --- | --- |
| *sampling time (pre vs. post)* | 1 | 13694 | 13694 | 4.4873 | 0.01651 | 0.001*** |
| *fuel manip. treatment* | 5 | 26096 | 5219.1 | 1.7102 | 0.03147 | 0.001*** |
| *pine proximity* | 1 | 6121 | 6120.5 | 2.0056 | 0.00738 | 0.001*** |
| *community type (litter vs. soil)* | 1 | 23908 | 23907.8 | 7.8342 | 0.02883 | 0.001*** |
| *fire unit* | 1 | 7110 | 7109.7 | 2.3297 | 0.00857 | 0.001*** |
| *time x fuel* | 4 | 15880 | 3970 | 1.3009 | 0.01915 | 0.001*** |
| *time x pine* | 1 | 4350 | 4349.5 | 1.4253 | 0.00524 | 0.001*** |
| *fuel x pine* | 5 | 20417 | 4083.5 | 1.3381 | 0.02462 | 0.001*** |
| *time x comm. type* | 1 | 8319 | 8319.2 | 2.726 | 0.01003 | 0.001*** |
| *fuel x comm. type* | 5 | 17786 | 3557.3 | 1.1657 | 0.02145 | 0.001*** |
| *pine x comm. type* | 1 | 3905 | 3905.4 | 1.2797 | 0.00471 | 0.005** |
| *patch within fire unit* | 19 | 88123 | 4638.1 | 1.5198 | 0.10626 | 0.001*** |
| *time x fuel x pine* | 4 | 14405 | 3601.2 | 1.1801 | 0.01737 | 0.002*** |
| *time x fuel x comm.* | 4 | 14052 | 3513.1 | 1.1512 | 0.01694 | 0.008** |
| *time x pine x comm.* | 1 | 3617 | 3617.4 | 1.1854 | 0.00436 | 0.038* |
| *fuel x pine x comm.* | 5 | 16265 | 3252.9 | 1.0659 | 0.01961 | 0.061 |
| *time x fuel x pine x comm.* | 4 | 14275 | 3568.7 | 1.1694 | 0.01721 | 0.009** |
| *residuals* | 174 | 531001 | 3051.7 |  | 0.64028 |  |

\*p < 0.05, \*\*p < 0.01, \*\*\* p < 0.0001

Shifts in fungal community composition corresponded to fire effects on fungal diversity and community heterogeneity. Increasing fuel loads reduced post-fire fungal diversity (inv. Simp. metric), and this effect was stronger on litter than soil fungi (F_5,144_=2.3, p=0.048; Fig. 4d,e, Table S15-16). Fire reduced community heterogeneity and had the strongest filtering effect in higher fuel loading treatments (F_5_,_243_=8, p<0.0001; Table S17). Fungi adapted to fire effects (Fox et al. 2022) were consistently found in post-fire plots, with indicator taxa varying with increasing fuel load and soil vs. litter location (Tables S18-S27). To summarize, changes in fungal community composition, including selection for fire-adapted taxa, were related to higher fuel consumption and soil heating, and associated with reductions in diversity and community heterogeneity, as well as selection for adaptation to surviving fire and in post-fire environments.

### 3.5. Fire slowed microbial decomposition

During the eight months following prescribed fire, microbial decomposition rates (k) were lower in burned relative to no burn plots (F_5_,_66_=2.16, p=0.069; Fig. S6, Table 6), with the greatest reductions in the 2X fuel addition plots. Decomposition rates linearly declined over time, but differences among fuel load treatments did not vary over time (Figure 7; F_15_,_66_=1.22, p=0.278). Despite similar effects of fire across time, post-hoc contrasts (Table S29) did suggest that fire effects on decomposition were strongest early after fire. In conclusion, plots that burned in 2017 had slower k-rates, but this effect did not vary across fuel load treatments.

**Table 6:** MANOVA results for fuel manipulation and pine proximity effects on microbial decomposition rate (k).

| model term | d.f. 1 | d.f. 2 | F-test | p-value |
| --- | --- | --- | --- | --- |
| *fuel manip. treatment* | 5 | 66 | 2.157 | 0.0694. |
| *pine proximity* | 1 | 66 | 0.011 | 0.9156 |
| *fire unit* | 1 | 66 | 6.532 | 0.0129 |
| *repeated measure* | 3 | 66 | 422.225 | <0.0001** |
| *fuel x pine* | 5 | 66 | 1.387 | 0.2407 |
| *fuel x rep. meas.* | 15 | 66 | 1.223 | 0.278 |
| *pine x rep. meas.* | 3 | 66 | 0.993 | 0.4015 |
| *unit x rep. meas.* | 3 | 66 | 2.373 | 0.0781 |
| *fuel x pine x rep. meas.* | 15 | 66 | 1.067 | 0.4032 |

. = p<0.1; \* = p<0.05; \*\* = p<0.0001

### 3.6. SEM – fuel manipulation effects on heat release and soil heating

Structural equation modeling synthesized how increasing fuels (and pine proximity) were linked to the system controlling fuel decomposition (Fig. 5, Table S30). For all SEM results, standardized regression coefficients (numbers in parenthesis) allow for comparison of pathways, and contrasting “direct” (d) effects between variables from “indirect” (i) effects, where another variable(s) mediates effects between variables. The model shows that adding fine fuels to plots (d: 0.889; Fig. S31) and being located near overstory pines (d: 0.363) increased soil heating, as described by peak surface and soil temperatures. This heating in turn drove greater fine fuel combustion (d: 0.662). Additionally, fine fuel combustion was lower near overstory pines due to unmeasured mechanisms (d: -0.391). The fuel manipulation treatments successfully increased soil heating temperature, duration, and drove differences in fine fuel combustion.

**Figure 5:**
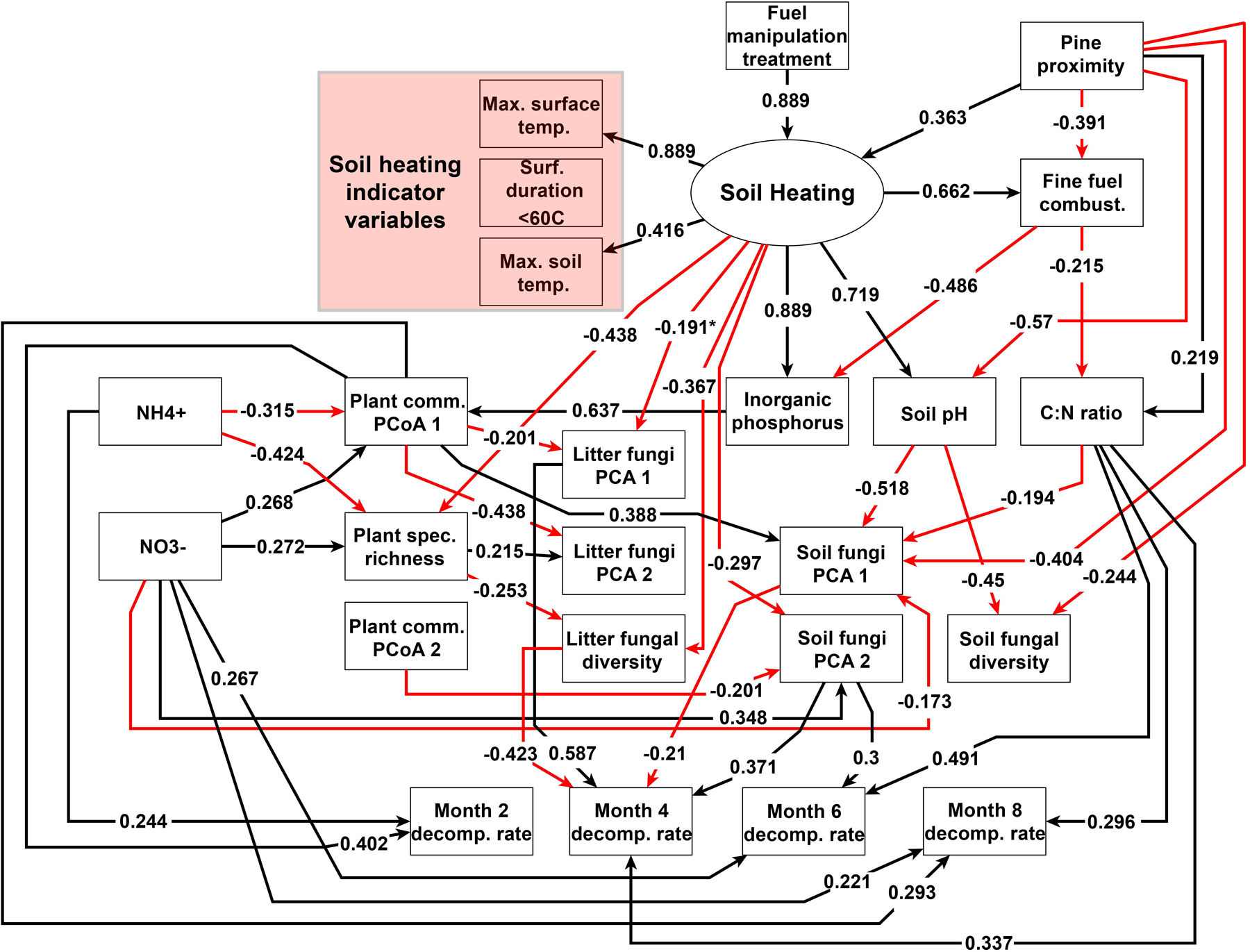
Final SEM for soil heating effects on microbial decomposition. Red and black paths represent negative and positive correlations between variables, respectively. Path coefficients are standardized regression coefficients, which allow for direct comparison of effects between model paths. Variables in the read box are the fire characteristics used as indicator variables for the soil heating latent variable.

### 3.7. SEM –soil heating effects on soils, plants, and fungi

As a product of fuel manipulation, soil heating and combustion were associated with changes in abiotic soil factors, plants, and fungi (Fig. 6). Greater soil heating drove direct increases in phosphorous (d: 0.555; Fig. 6a) that complimented greater phosphorous levels near pines (i: 0.158); however, these increases were partially offset due to increased fuel combustion associated with soil heating (i: -0.322). Soil heating also caused indirect decreases in C:N ratios through fuel combustion (i: -0.142), and this effect was only weakly modified by pine proximity (i: -0.052). Greater soil heating also drove direct increases in soil pH (d: 0.719) that negated pine proximity effects on soil pH (lower near pines; d: -0.256). Soil heating did not influence ammonium (NH_4_ ^+^) or nitrate (NO ^-^) availability.

**Figure 6:**
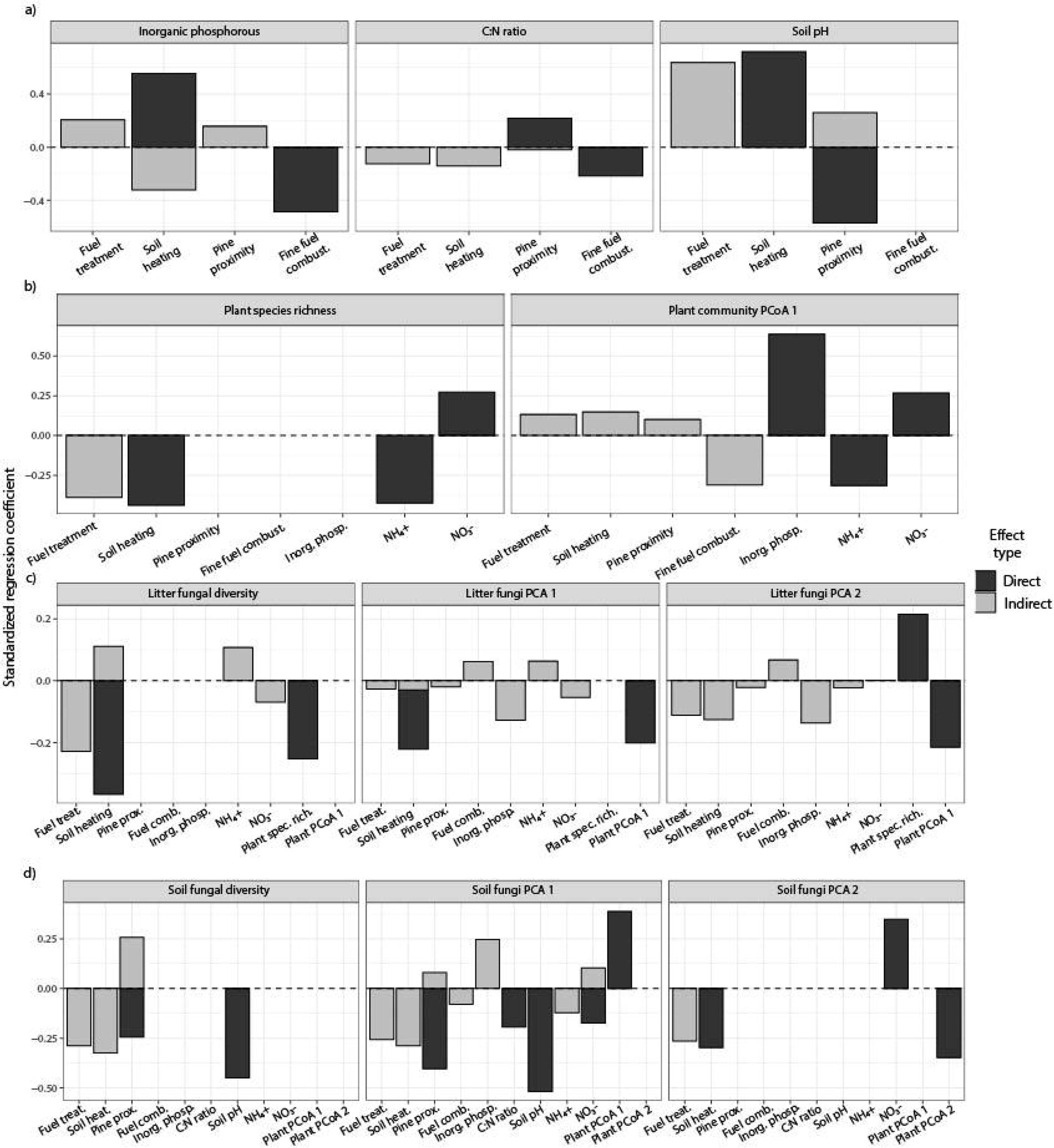
SEM results - Soil heating effects on soils, plants, fungi, and their interactions. Direct effects (black) represent paths linking the response and independent variable, while indirect effects (grey) represent paths between variables that are mediated by one or more variables. Soil heating altered a) soil chemistry through fine fuel combustion, and b) plant communities by reducing species richness and altering P and N availability. Fire c) directly altered litter fungal communities, as well as through changes to plant community composition. Fire effects on d) soil fungal communities were mediated by changes to soil pH, nutrient availability, and plant community composition.

Plant community diversity and composition were governed by soil heating and soil characteristics (Fig. 6b). Both increases in soil heating (d: -0.438) and ammonium (d: -0.424) were correlated with reduced plant species richness, but higher nitrate availability was linked to higher richness values (d: 0.272). Plant community composition was primarily controlled by nutrient availability, with P being most important (d: 0.637), followed by ammonium (d: -0.315), and nitrate (d: 0.268). Soil heating also had a small, indirect effect on plant community composition through changes to P availability (i: 0.148).

Fungal communities were altered by soil heating but the community types responded differently to plant community composition and soil factors (Fig. 6c, d). Soil heating reduced both litter (d: -0.367) and soil (i: -0.324) fungal diversity. Declines in soil fungal diversity were mediated by increases in soil pH associated with soil heating (d: -0.518) while litter fungal diversity was inversely related to plant community diversity (d: -0.253). The composition of litter fungal communities was influenced by soil heating (d: -0.191; i: -0.126), plant community composition (d: -0.201, -0.215) and plant species richness (d: 0.215). Soil fungal composition was also directly influenced by soil heating (d: -0.297) and soil heating effects on soil pH (i: - 0.372). However, soil pH (d: -0.518), pine proximity (d: -0.404), nitrate (d: 0.348), and plant community composition (d: 0.388, -0.348) were the strongest predictors of soil fungal composition. To conclude, soil heating drove changes to abiotic soil factors and plant communities that influenced fungal community composition.

### 3.8. SEM – soil heating effects on decomposition

Fuel loading effects on soil heating had specific direct and indirect effects on decomposition that varied in the 8 months after fire (Fig. 7; Table S33). Two months after fire, plant composition (d: 0.402, via soil heating effects on P availability) and a fire-independent effect of ammonium (d: 0.244) were the primary determinants of decomposition. Four and six months after fire, soil heating driven changes to soil C:N ratios (d: 0.337, 0.491) and litter (d: -0.423, 0.587) and soil (d: -0.21, 0.371, 0.3) fungal communities and diversity (d: -0.423) were the most important drivers of decomposition. Eight months after fire, soil heating related changes to plant composition (d: 0.293) and soil C:N ratios (d: 0.296), as well as a fire independent effect of nitrate (d: 0.221) governed decomposition. To summarize, soil heating effects on soil abiotic, plant, and fungal components slowed the decomposition of new fuels throughout the eight months following fire.

**Figure 7:**
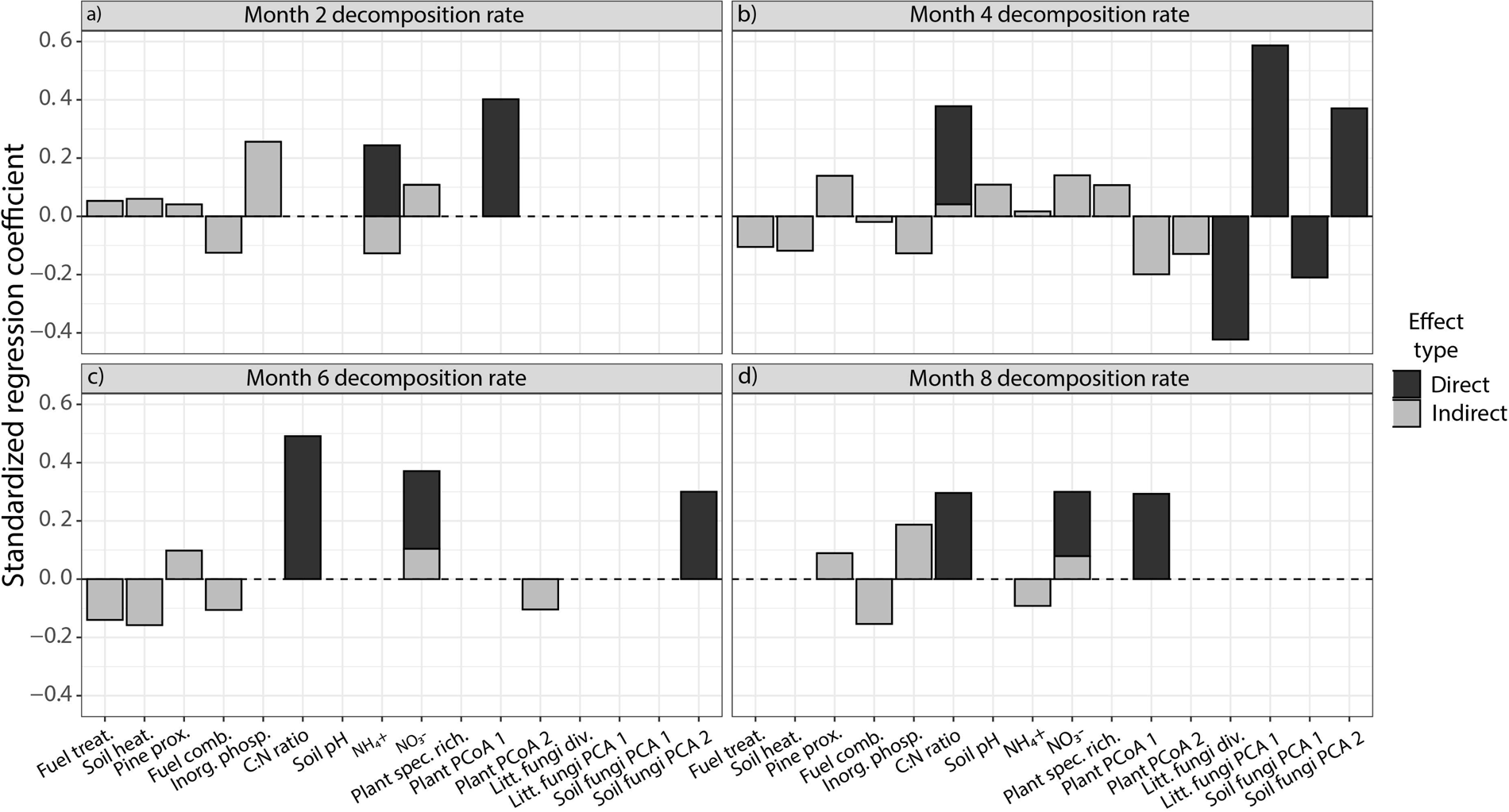
SEM results - Soil heating and environmental effects on decomposition (k). a) Two months after fire, ammonium and plant community composition (influenced by soil heating through changes to P availability) were the dominant predictors on k-rate. b) Four months after fire, fungal community composition (shifted with soil heating) and C:N ratios (decreased with soil heating) had the strongest effect on k-rates. C) 6 months post-fire, C:N ratios, and soil fungal community composition had the strongest influence on decomposition. d) Eight months post-fire, C:N ratios, ammonium, and plant community composition determined k-rates.

## 4. Discussion

### In our study, increased fuel loads made fires hotter and longer, which reduced post-fire decomposition of new fuels

This slower decomposition was mediated by soil heating effects on plant communities and soil abiotic & biotic components, and these effects persisted over the eight months after fire. Fuel buildup that creates more intense fires is well known to impact vegetation recovery in ecosystems with prescribed burning and wildfires (Stephens et al., 2009; Volkova et al., 2019). Fuel reduction is a key management tool to reduce wildfire impacts in fire-suppressed systems (Schwilk et al., 2009; Stephens et al., 2009). Our results show fuel loads also weigh on postfire decomposition of new fuels. Eight months after fire, the quantity of new fuels decomposed in plots with larger fuel loads (add 1X-2X) were 25-32% lower than unburned plots and 10-18% lower than fuel removal plots. These effects may be applicable across systems.

Warm and wet pyrophilic ecosystems like savannas may be more likely to show post-fire differences based on fuel load because of rapid decomposition and high productivity, compared to colder, drier ecosystems (Archibald et al., 2018). However, many drier systems are historically fire suppressed, so differences in decomposer responses to fire, including their ability to breakdown new fuels, may be even greater than those with fire-adapted decomposers and vegetation substrates. In past work, post-fire fuel accumulation is nearly always attributed to vegetation responses alone and varies widely based on climate (Coppoletta et al., 2016; Tepley et al., 2018; Volkova et al., 2019), but these results show that decomposition response to fire may play an equally important role to fire-fuel feedbacks.

### Decomposition shifts in response to increased soil heating were complex but tied to immediate postfire changes in abiotic and biotic properties

Due to logistical limitations, we could not track many ecosystem properties over the year following fire, but immediate fire responses of the system still helped predict decomposition shifts. Litter fungi were more impacted by fire than soil fungi, especially with large fuel additions, which supports an insulating role for soil shown in previous findings (Dao et al., 2022; SemenovaLJNelsen et al., 2019). These rapid (∼1-2 months) fungal responses to fire were directly tied to changes in decomposition much later, at 4 and 6 months after fire (Fig. 5). More intense fires have greater impacts on fungi (Day et al., 2019; Holden et al., 2016; Mino et al., 2021) in ways that more drastically alter postfire fungal succession and their functional ability in the growing season after fire. Adding fuels also caused increasingly greater (albeit modest) shifts to the fire-adapted plant communities (Fig 3) in this system (Beckage et al., 2009), similar to other “pyrophilic” ecosystems (Archibald et al., 2018). These plant responses to fire influenced decomposition early and late in the season (Fig. 5), likely as a product of soil moisture and priming by plant exudates (Cheng, 2009; Yan et al., 2023). However, a follow-up survey of plants in the summer of 2018 found no differences among fuel-manipulation plots, showing the resilience of these fire-adapted plant communities. Interestingly soil carbon and most nitrogen components did not respond to hotter fires, even when pH, P, and ammonium did. Hotter fires should have volatilized more N and thereby altered C:N ratios (Certini, 2005; Raison, 1979), critical forces shown to influence decomposition in previous work (Butler et al., 2019; Cornelissen et al., 2017; Ficken and Wright, 2017; Hopkins et al., 2020; SemenovaLJNelsen et al., 2019). In our study, these nutrients were still key controls on decomposition, despite their resilience to more intense fires, with greater post-fire nitrate levels and C:N ratios linked to faster microbial decomposition of new fuels.

Combined, these interacting factors show that, by altering fire characteristics, variation in fuels can influence the ecosystem components that govern the breakdown of new fuels available to future fires.

### Natural variation in fuel loads on the landscape is likely to produce pyrodiversity not just in organisms but also their ecological functions

We experimentally manipulated fuel loads both near and away from canopy pines to levels likely to occur as patches on the landscape. Pine needle drops from winds and storms can easily create the 2X conditions that produced the hottest fires, whereas local topography can often produce patches that fire doesn’t reach. Just as experimental variation in fuels differentially impacted plants, fungi, and soil properties, natural fuel variation on the landscape should produce a patchwork of niches and biota. More than a dozen fungi were restricted to pre-burn communities, including several saprotrophic and plant associated fungi, while post-fire communities were dominated by black yeasts, plant pathogens, and mycoparasites which is consistent with other systems (Fox et al., 2022; Hopkins et al., 2021). Pre and postburn indicator taxa also differed depending on fuel treatment, a reflection that fuel variation is an important component of the pyrodiversity-biodiversity link for fungi (Jones and Tingley, 2021). As the dominant plant mutualists, pathogens, and decomposers of plant fuels in this system, shifts in fungi differentially impact new fuel production and decomposition and, on a landscape scale, their combined forces can help maintain a dynamic landscape of fuels that reinforce pyrodiversity (Fox et al., 2022; Jones and Tingley, 2021). Many of these dynamics take place over multiple years and so directly link to the resilience of pyrophilic ecosystems, many of which are threatened and face increasing risks with a changing climate.

### For the restoration and conservation of “pyrophilic ecosystems” like savannas and grasslands, fuel management is a consistent tool that can help reinforce fire feedbacks through plant and soil systems

A key aspect of our work is that a heterogenous distribution of fuels across the landscape clearly helps drive pyrodiversity, including fire effects on plant and fungal biodiversity, soil properties, and ultimately the breakdown of new fuels on which future fires will rely. Fire-suppression across huge management units has built up abundant, continuous fuels and supported vegetation (fuel producers) and microorganisms, like fungi (fuel decomposers) that may be otherwise rare and localized in systems with recurrent fire (Holzmueller et al., 2008; Olsson and Jonsson, 2010; Stephens et al., 2009; Volkova et al., 2019). In the same way, the reintroduction of fire without fuel management, whether intentional or through wildfire, is unlikely to create the pyrodiversity that makes these systems resilient. In areas with large fuel loads, pre-fire work to create a patchy distribution of fuels on the landscape (Fiedler et al., 1998; Stephens et al., 2021) can create a variety of environments and niches and support the biotic functions that feedback to fuels. Taking advantage of existing heterogeneity in fuel production, such as proximity to overstory trees, can aid fuel feedbacks to support pyrodiversity and system resilience. The explicit incorporation of decomposition responses to fire in relation to fuel management will be sorely needed as fire and the many components that build and breakdown fuels shift with climate (Jones et al., 2022; Moritz et al., 2012). Since more than 40% of Earth’s terrestrial ecosystems depend on recurrent fires like these, their conservation relies on greater work to understand these complex and interacting responses to fire.

## Supporting information

Full Supplemental Information

## Acknowledgements

The authors would like to thank Sam Imel, Joshua Faylo, Paige Hansen, Juna Murao, Hannah Dea, Dr. Donald Nelsen, Tall Timbers Research Station and Land Conservancy, and the Wade Foundation for lab assistance and field support. We also thank Paddy Wade for providing access to the property, Cinnamon Dixon, Monica Rother, and Michelle Smith for mapping burned areas, and Eric Staller, Andrew Chase, and other Tall Timbers and Arcadia Plantation staff for assisting with prescribed fires. The authors also thank Dr. Candace Davison at the Pennsylvania State Breazeale Nuclear Reactor for assistance with gamma sterilization of plant litter.

This work was supported by an NSF Graduate Research Fellowship (1540502) to JRH, as well as NSF grants DEB-1557000 to BAS and DEB-1556837 to WJP.

## Competing interests

The authors have no competing interests to declare.

## Author contributions

WJP, BAS, and JMH conceived and designed the experiment. JRH, JMH, NJJ, TAS, WJP, and BAS performed the experiment. KMR managed the March and April 2017 prescribed fires. TAS prepared the fungal ITS libraries. JRH analyzed the data and wrote the manuscript. JMH, NJJ, TAS, WJP, KMR, and BAS provided editorial advice.

## Data availability

Upon publication, sequence data will uploaded to the NCBI database and all other data will be uploaded to Dryad.

## Notes

### Competing Interest Statement

The authors have declared no competing interest.

