## Supplementary material for "Fuel buildup shapes post-fire fuel decomposition through soil heating effects on plants, fungi, and soil chemistry": Full Supplemental Information

**New Phytologist Supporting Information**

**Article acceptance date:**

**Contents:**

Section 1: Wade Tract site map – pg. 2

Section 2: Fuel load manipulation/soil heating treatments – pg. 3

Section 3: Structural equation modeling procedures – pg. 5

Section 4: Fuel manipulation effects on fire characteristics – pg. 14

Section 5: Soil heating effects on abiotic soil factors – pg. 18

Section 6: Soil heating effects on plant communities – pg. 22

Section 7: Soil heating effects on fungal communities – pg. 24

Section 8: Soil heating effects on microbial decomposition – pg. 39

Section 9: Structural equation model results – pg. 43

References – pg. 47

**Appendix Section 1: Wade Tract Site Map**

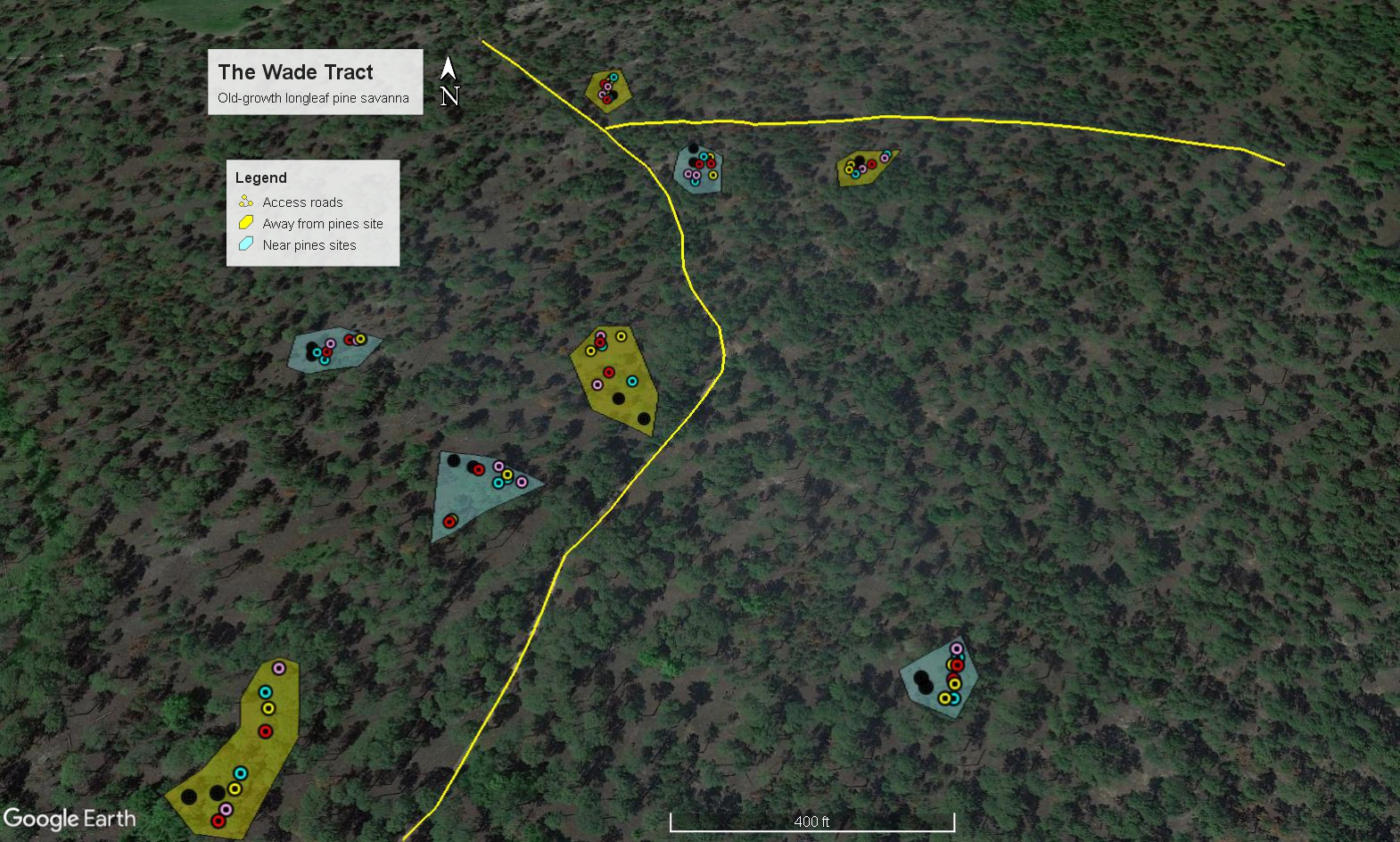

**Figure S1:** The Wade Tract Site Map. Yellow lines represent access roads, yellow polygons are away from pines patches, and blue polygons are near pines patches. Points represent fuel manipulation plots: white = no burn, light blue = fuel removal, black = reference, yellow = switch, light pink = add 1X pine needles, and red = add 2X pine needles. No burn plots do not always appear in patches as they had to be located following 2017 prescribed fires.

**Appendix Section 2: Fuel load manipulation/soil heating treatments**

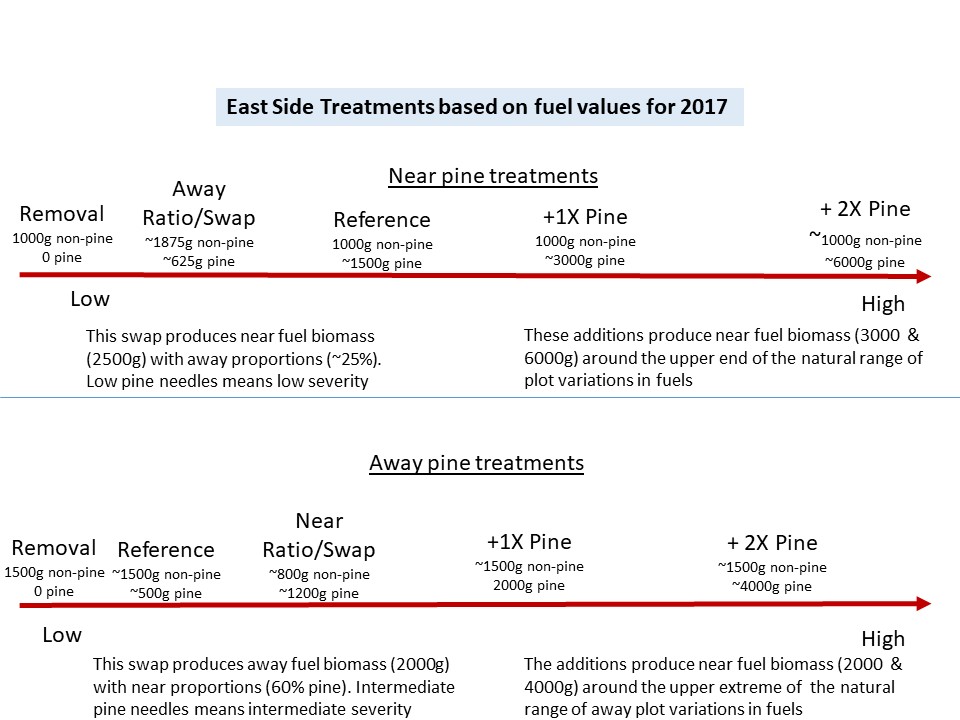

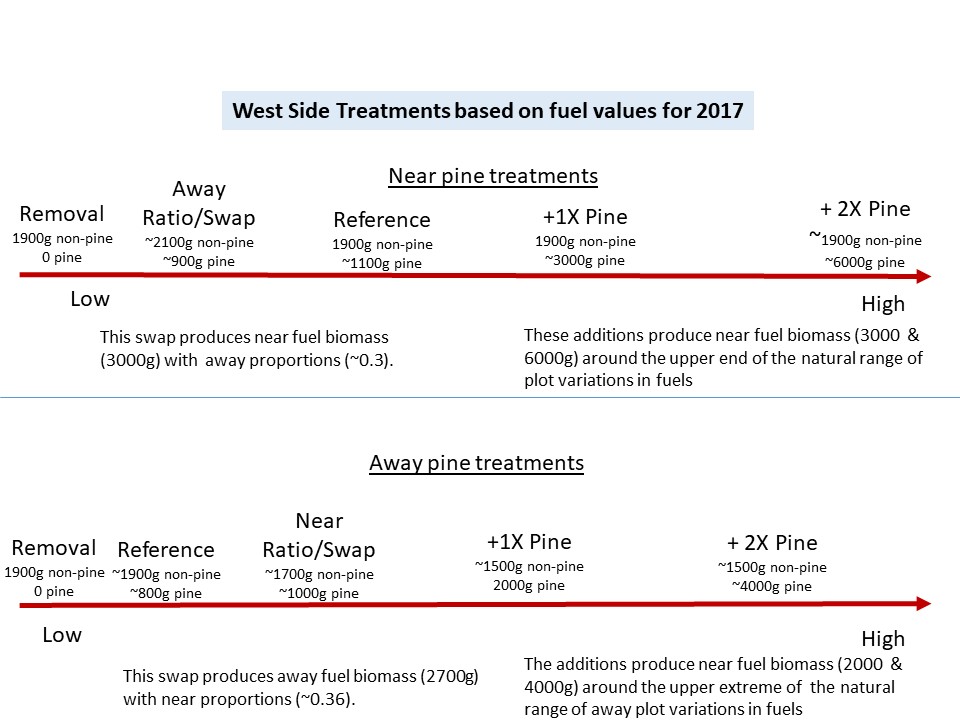

**Figure S2:** Fuel load manipulations for each fire unit. These treatments were designed to create a similar range of soil heating near and away from pines. Different amounts of pine needles were applied in near and away plots to produce different fuel loads ranging from 0-1000 g/4m^2^. The addition plots treatments add the same amounts in both near and away to generate the amount of pine needles above the mean amounts in near plots.

**Section 3: Structural equation modeling procedures**

**Table S3:** Initial SEM framework. The design of the initial SEM is described below. Note that citations relate to expected hypotheses based on the scientific literature.

| **Response Variable** | **Explanatory Variable(s)** | **Citation** |
| --- | --- | --- |
| soil heating (latent variable) | Max. surf. temp. change |  |
|  | Surf. dur. >60C |  |
|  | Max. soil temp. change |  |
| soil heating | fuel load treatment | (Ellair & Platt, 2013) |
|  | pine proximity | (Platt *et al.*, 2016) |
| fine fuel combustion | soil heating |  |
|  | pine proximity | (Platt *et al.*, 2016) |
| fungal pca axis 1 & 2 (litter and soil) | soil heating | (Bárcenas-Moreno & Bååth, 2009) |
|  | pine proximity | (Semenova‐Nelsen *et al.*, 2019) |
|  | plant pcoa axis 1 | (Carson *et al.*, 2019) |
|  | plant pcoa axis 2 | (Carson et al. 2019) |
|  | plant species richness | (Carson et al. 2019) |
|  | inorg. phosp. | (Tedersoo *et al.*, 2014) |
|  | C:N ratio | (Lauber *et al.*, 2008) |
|  | soil pH | (Tedersoo *et al.,* 2014) |
|  | NH4+ | (Tedersoo *et al*., 2014) |
|  | NO3- | (Tedersoo *et al.,* 2014) |
| fungal inverse simpson index (litter and soil) | soil heating |  |
|  | pine proximity | (Semenova-Nelsen *et al.* 2019 |
|  | plant pcoa axis 1 | (Shen *et al.*, 2021) |
|  | plant pcoa axis 2 | (Shen *et al.*, 2021) |
|  | plant species richness | (Shen *et al.*, 2021) |
|  | inorg. phosp. | (Tedersoo *et al.,* 2014) |
|  | C:N ratio | (Lauber *et al.*, 2008) |
|  | soil pH | (Tedersoo *et al.,* 2014) |
|  | NH4+ | (Tedersoo *et al.,* 2014) |
|  | NO3- | (Tedersoo *et al.,* 2014) |
| plant pcoa axis 1 and 2 | soil heating | (Gagnon *et al.*, 2015) |
|  | pine proximity | (Mugnani *et al.*, 2019) |
|  | inorg. phosp. | (Janssens *et al.*, 1998) |
|  | C:N ratio | (De Deyn *et al.*, 2008) |
|  | soil pH | (Jenks & Hasegawa, 2008) |
|  | NH4+ | (De Deyn *et al.*, 2008) |
|  | NO3- | (De Deyn *et al.*, 2008) |
| plant comm. richness | soil heating | (Grace & Keeley, 2006) |
|  | pine proximity | (Mugnani *et al.*, 2019) |
|  | inorg. phosp. | (Janssens *et al.*, 1998) |
|  | C:N ratio | (De Deyn *et al.*, 2008) |
|  | soil pH | (Jenks & Hasegawa, 2008) |
|  | NH4+ | (De Deyn *et al.*, 2008) |
|  | NO3- | (De Deyn *et al.*, 2008) |
| inorg. phosp. | soil heating | (Butler *et al.*, 2018) |
|  | pine proximity |  |
|  | fine fuel combustion | (Butler *et al.*, 2018) |
| NH_4_^+^ | soil heating | (Raison, 1979) |
|  | pine proximity | (Perry *et al.*, 2008) |
|  | fine fuel combustion | (Johnson & Curtis, 2001) |
| NO_3_^-^ | soil heating | (Raison, 1979) |
|  | pine proximity | (Perry *et al.*, 2008) |
|  | fine fuel combustion | (Johnson & Curtis, 2001) |
| C:N ratio | soil heating | (Johnson & Curtis, 2001) |
|  | pine proximity | (Perry *et al.*, 2008) |
|  | fine fuel combustion | (Johnson & Curtis, 2001) |
| soil pH | soil heating | (Certini, 2005) |
|  | pine proximity |  |
|  | fine fuel combustion | (Certini, 2005) |
| decomposition rate K (months 2, 4, 6, and 8) | soil heating | (Bárcenas-Moreno & Bååth, 2009) |
|  | fungal pca axis 1 | (van der Wal *et al.*, 2013) |
|  | fungal pca axis 2 | (van der Wal *et al.*, 2013) |
|  | fungal inverse simpson index | (van der Wal *et al.*, 2013) |
|  | plant pcoa axis 1 | (Cheng *et al.*, 2003) |
|  | plant pcoa axis 2 | (Cheng *et al.*, 2003) |
|  | plant comm. richness | (Cheng *et al.*, 2003) |
|  | inorg. phosp. | (Butler *et al.*, 2019) |
|  | C:N ratio | (Manzoni *et al.*, 2010) |
|  | NH4+ | (Manzoni *et al.*, 2010) |
|  | NO3- | (Manzoni *et al.*, 2010) |
|  | soil pH | (Leifeld *et al.*, 2008) |

**Table S4:** Model fit statistics for SEMs. RMSEA = root mean square error (p > 0.05 implies good fit). CFI = comparative fit index (p > 0.97 implies good fit). RMR = root mean square residual (0-0.1 is adequate). NNFI = non-normed fit index (> 0.98 is a good fit). SRMR = square root mean square (0-0.08 is a good fit).

| Model | p-value | RMSEA 90% CI | RMSEA CI p-value | CFI | RMR | NNFI | SRMR | AIC | BIC |
| --- | --- | --- | --- | --- | --- | --- | --- | --- | --- |
| 1 | 0 | 0.145-0.189 | 0 | 0.686 | 0.126 | 0.221 | 0.106 | 3738.515 | 4142.951 |
| 2 | 0.004 | 0.047-0.11 | 0.063 | 0.935 | 0.059 | 0.818 | 0.06 | 3591.559 | 4024.262 |
| 3 | 0.13 | 0-0.08 | 0.532 | 0.972 | 0.06 | 0.938 | 0.061 | 3542.805 | 3921.148 |
| 4 | 0.364 | 0-0.063 | 0.832 | 0.991 | 0.073 | 0.985 | 0.072 | 3500.489 | 3813.601 |
| 5 | 0.503 | 0-0.054 | 0.921 | 1 | 0.088 | 1.002 | 0.086 | 3284 | 3531.88 |

Final structural equation model (model 5)

mod_all5 <- '

\# soil heating variable

intensity =~ SurfPeak + SurfDurSec + SoilPeak

\# regression paths

intensity ~ sev.tre + pine.code

combustion ~ intensity + pine.code

\# fungi

litt_pca1 ~ intensity + plant.pcoa1 + pH + plant.pcoa2

litt_pca2 ~ plant.pcoa1 + plant.rich + cn + nh4 + no3

litt_comp.invsimp ~ intensity + plant.rich + cn

soil_pca1 ~ pine.code + plant.pcoa1 + cn + pH + no3

soil_pca2 ~ intensity + plant.pcoa2 + cn + no3

soil_comp.invsimp ~ pine.code + plant.pcoa2 + totp + pH + nh4

\# plants

plant.pcoa1 ~ totp + nh4 + no3

plant.rich ~ intensity + pine.code + totp + pH + nh4 + no3

\# soils

totp ~ intensity + pine.code + combustion

cn ~ pine.code + combustion

pH ~ intensity + pine.code

\# decomp

krate2 ~ litt_pca1 + plant.pcoa1 + totp + cn + pH + nh4

krate4 ~ soil_pca1 + soil_pca2 + litt_pca1 + litt_comp.invsimp + plant.pcoa1 + plant.pcoa2 + cn

krate6 ~ soil_pca1 + soil_pca2 + soil_comp.invsimp + litt_pca2 + plant.pcoa2 + cn + nh4 + no3

krate8 ~ litt_pca1 + plant.pcoa1 + plant.pcoa2 + totp + cn + no3

\# covariance paths

soil_comp.invsimp ~~ soil_pca1 + soil_pca2

litt_comp.invsimp ~~ litt_pca1 + litt_pca2

litt_pca1 ~~ litt_pca2

plant.rich ~~ plant.pcoa1

soil_pca1 ~~ soil_pca2

cn ~~ totp + nh4 + no3 + pH

'

**Section 4: Fuel manipulation effects on fire characteristics**

**Table S5:** Contrasts for fire characteristic and fuel combustion MANOVA model.

| contrast | estimate | s.e. | d.f. | t-ratio | p-value |
| --- | --- | --- | --- | --- | --- |
| *away vs. near* | 7.15 | 5.36 | 66 | 1.334 | 0.1868 |
| *removal vs. reference* | -13.68 | 3.29 | 66 | -4.155 | 0.0001*** |
| *removal vs. switch* | -9.19 | 3.53 | 66 | -2.601 | 0.0115* |
| *removal vs. 1x* | -20.64 | 3.29 | 66 | -6.271 | <0.0001*** |
| *removal vs. 2x* | -22.59 | 3.29 | 66 | -6.864 | <0.0001*** |
| *reference vs. switch* | 4.49 | 3.53 | 66 | 1.271 | 0.208 |
| *reference vs. 1x* | -6.97 | 3.29 | 66 | -2.116 | 0.0381* |
| *reference vs. 2x* | -8.91 | 3.29 | 66 | -2.709 | 0.0086* |
| *switch vs. 1x* | -11.46 | 3.53 | 66 | -3.244 | 0.0019* |
| *switch vs. 2x* | -13.4 | 3.53 | 66 | -3.796 | 0.0003** |
| *1x vs. 2x* | -1.95 | 3.29 | 66 | -0.592 | 0.5557 |
| . = p<0.1; * = p<0.05; ** = p<0.0001 | | | | | |

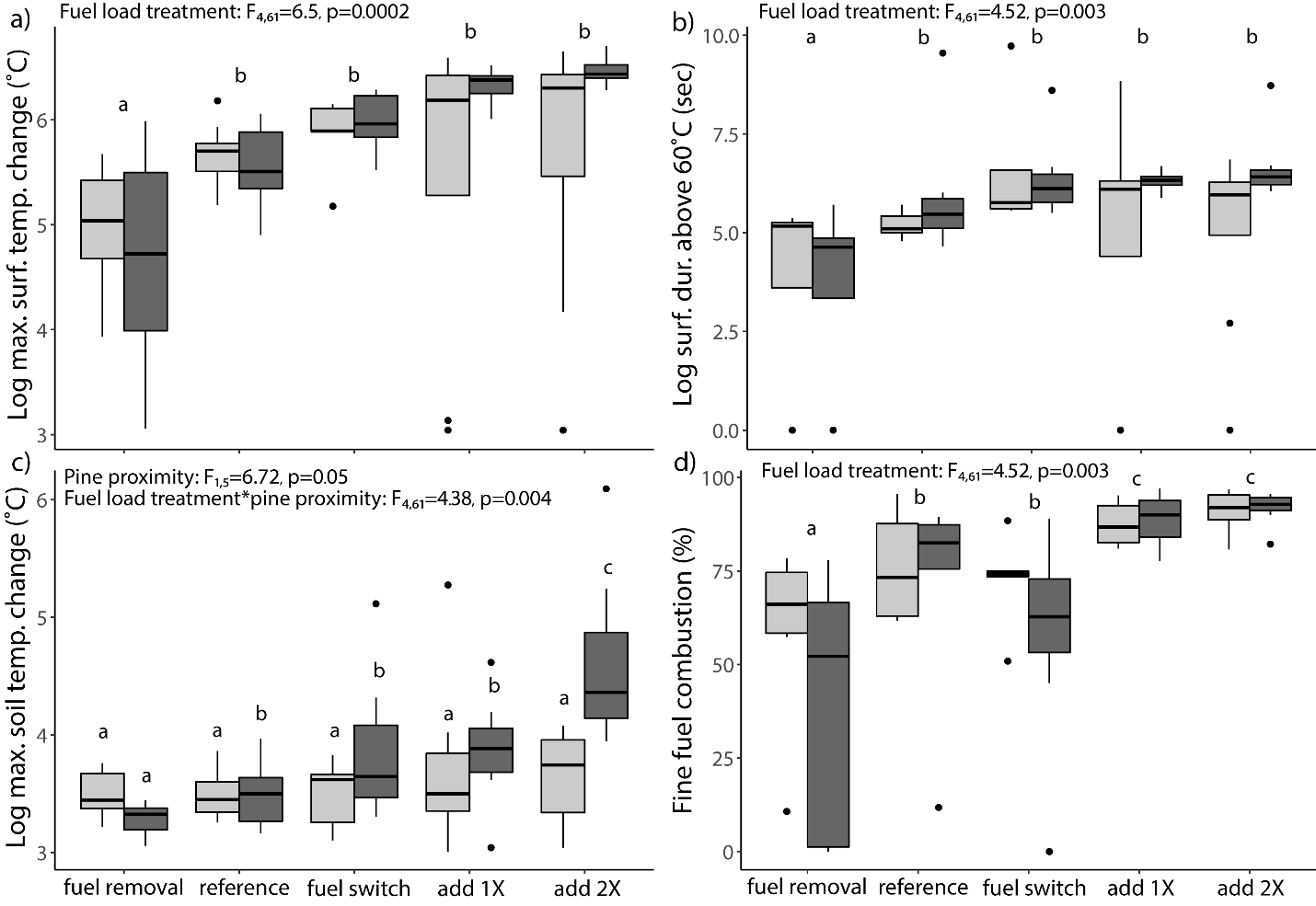

**Figure S6:** Fuel manipulation and pine proximity effects on soil heating and fine fuel combustion. Light grey and dark grey boxes represent “away from pines” and “near pines” plots, respectively. Lower case letters denote significant differences (p<0.05) at the fuel load treatment level. As fuel manipulations added more fuels, a) maximum changes in surface temperature, b) surface fire durations >60°C, c) maximum changes in soil temperature, and d) fine fuel combustion increased.

**Section 5: Soil heating effects on abiotic soil factors**

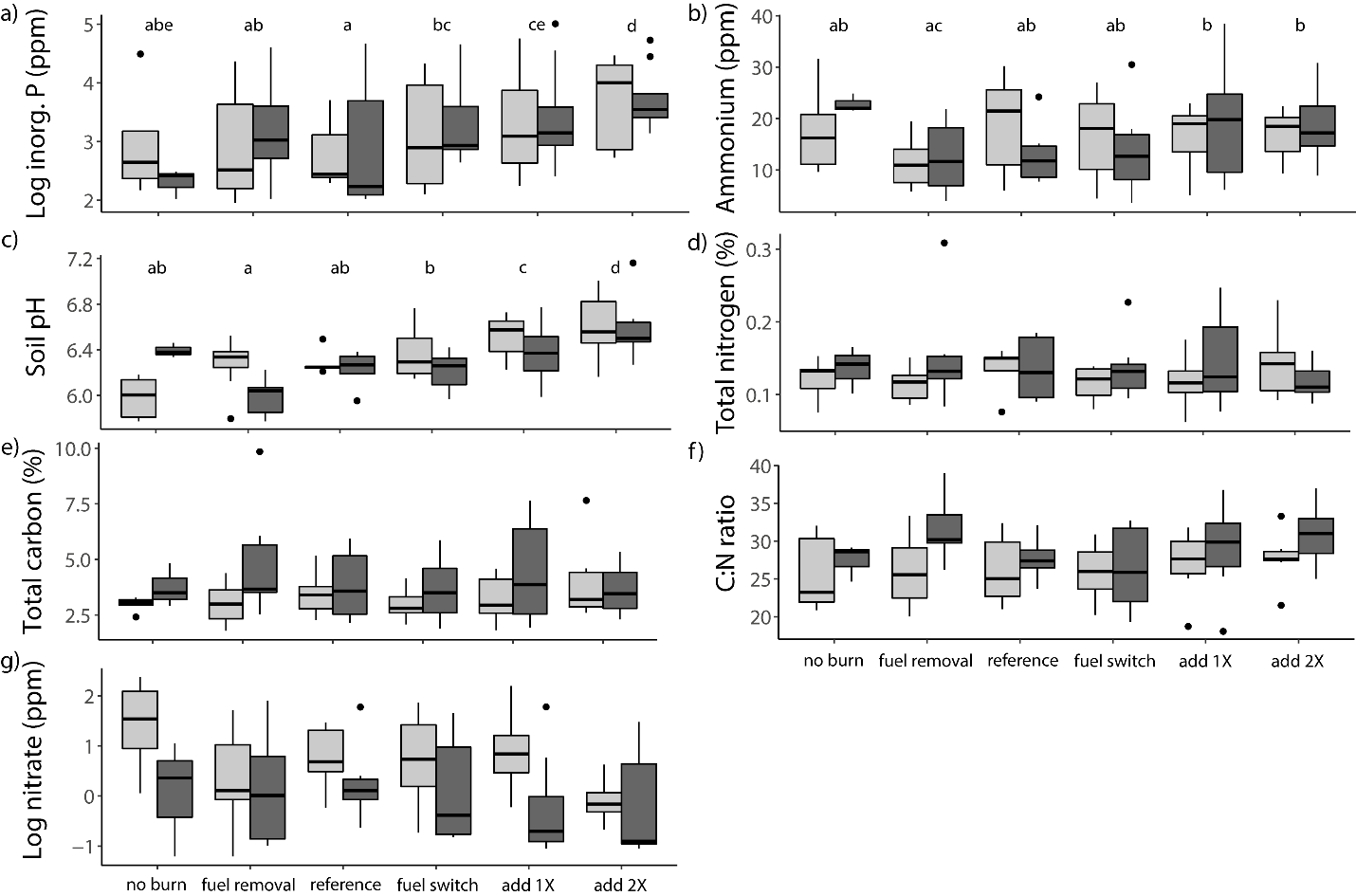

**Figure S7:** Fuel manipulation and pine proximity effects on soil factors. Light grey and dark grey boxes represent “away from pines” and “near pines” plots, respectively. Lower case letters denote significant differences (p<0.05) among fuel load treatments only. As soil heating increased, a) inorganic phosphorus, b) ammonium, c) and soil pH levels increased. Fire severity and pine proximity did not have a strong influence on d) total nitrogen, e) total carbon, f) C:N ratios, or g) nitrate levels.

**Table S8:** Contrasts for inorganic phosphorus and ammonium models.

|  | Inorg. Phosph. | | | Ammonium | | |
| --- | --- | --- | --- | --- | --- | --- |
| contrast | estimate | t-ratio | p-value | estimate | t-ratio | p-value |
| *away vs. near* | 0.368 | 0.141 | 0.8907 | -5.014 | -0.2 | 0.8461 |
| *no burn vs. removal* | -0.581 | -0.75 | 0.4679 | 15.683 | 1.966 | 0.0693 |
| *no burn vs. reference* | -0.248 | -0.318 | 0.7561 | 9.497 | 1.172 | 0.2596 |
| *no burn vs. switch* | -0.975 | -1.258 | 0.2323 | 9.393 | 1.174 | 0.2596 |
| *no burn vs. 1x* | -1.28 | -1.656 | 0.1238 | 5.322 | 0.67 | 0.514 |
| *no burn vs. 2x* | -1.806 | -2.332 | 0.0379* | 3.911 | 0.49 | 0.6315 |
| *removal vs. reference* | 0.333 | 1.364 | 0.1781 | -6.185 | -1.671 | 0.1003 |
| *removal vs. switch* | -0.395 | -1.737 | 0.088 | -6.289 | -1.824 | 0.0735 |
| *removal vs. 1x* | -0.7 | -3.182 | 0.0024* | -10.36 | -3.106 | 0.003** |
| *removal vs. 2x* | -1.226 | -5.472 | <0.0001*** | -11.771 | -3.464 | 0.001** |
| *reference vs. switch* | -0.727 | -2.941 | 0.0048** | -0.104 | -0.028 | 0.978 |
| *reference vs. 1x* | -1.032 | -4.29 | 0.0001*** | -4.175 | -1.143 | 0.2578 |
| *reference vs. 2x* | -1.558 | -6.383 | <0.0001*** | -5.586 | -1.508 | 0.1373 |
| *switch vs. 1x* | -0.305 | -1.362 | 0.1789 | -4.071 | -1.198 | 0.2361 |
| *switch vs. 2x* | -0.831 | -3.639 | 0.0006** | -5.482 | -1.582 | 0.1193 |
| *1x vs. 2x* | -0.526 | -2.393 | 0.0202* | -1.411 | -0.423 | 0.6739 |
| *p < 0.05, **p < 0.01, *** p < 0.0001 | | | | |  |  |

**Table S9:** Contrasts for soil pH model.

| contrast | estimate | t-ratio | p-value |
| --- | --- | --- | --- |
| *away vs. near* | 0.2792 | 0.916 | 0.3833 |
| *no burn vs. removal* | 0.116 | 0.611 | 0.5434 |
| *no burn vs. reference* | -0.1494 | -0.744 | 0.4595 |
| *no burn vs. switch* | -0.2115 | -1.104 | 0.2736 |
| *no burn vs. 1x* | -0.521 | -2.786 | 0.0071** |
| *no burn vs. 2x* | -0.8194 | -4.316 | 0.0001*** |
| *removal vs. reference* | -0.2653 | -1.569 | 0.1219 |
| *removal vs. switch* | -0.3275 | -2.084 | 0.0414* |
| *removal vs. 1x* | -0.637 | -4.191 | 0.0001*** |
| *removal vs. 2x* | -0.9354 | -6.047 | <0.0001*** |
| *reference vs. switch* | -0.0622 | -0.363 | 0.7175 |
| *reference vs. 1x* | -0.3717 | -2.232 | 0.0293* |
| *reference vs. 2x* | -0.67 | -3.959 | 0.0002*** |
| *switch vs. 1x* | -0.3095 | -2.001 | 0.0499* |
| *switch vs. 2x* | -0.6078 | -3.856 | 0.0003*** |
| *1x vs. 2x* | -0.2983 | -1.962 | 0.0543* |
| Away vs. near treatment effects | | | |
| *no burn* | -0.4151 | -2.583 | 0.012* |
| *removal* | -0.121 | -0.842 | 0.4029 |
| *reference* | 0.3045 | 2.436 | 0.0179* |
| *switch* | 0.1366 | 1.156 | 0.2528 |
| *1x* | 0.3128 | 2.855 | 0.0061* |
| *2x* | 0.2463 | 2.248 | 0.0288* |
| Away from pines treatment effects | | | |
| *no burn vs. removal* | -0.2941 | -2.376 | 0.0205* |
| *no burn vs. reference* | -0.3095 | -2.284 | 0.0257* |
| *no burn vs. switch* | -0.3915 | -3.11 | 0.0028** |
| *no burn vs. 1x* | -0.5479 | -4.427 | <0.0001*** |
| *no burn vs. 2x* | -0.6345 | -4.929 | <0.0001*** |
| *removal vs. reference* | -0.0154 | -0.127 | 0.8991 |
| *removal vs. switch* | -0.0974 | -0.889 | 0.3773 |
| *removal vs. 1x* | -0.2537 | -2.407 | 0.0192* |
| *removal vs. 2x* | -0.3404 | -3.107 | 0.0029** |
| *reference vs. switch* | -0.082 | -0.661 | 0.5108 |
| *reference vs. 1x* | -0.2383 | -1.97 | 0.0534* |
| *reference vs. 2x* | -0.325 | -2.605 | 0.0115* |
| *switch vs. 1x* | -0.1563 | -1.427 | 0.1587 |
| *switch vs. 2x* | -0.243 | -2.131 | 0.037* |
| *1x vs. 2x* | -0.0867 | -0.791 | 0.4319 |
| Near pines treatment effects | | | |
| *no burn vs. removal* | 0.4101 | 2.808 | 0.0066** |
| *no burn vs. reference* | 0.1602 | 1.065 | 0.2908 |
| *no burn vs. switch* | 0.18 | 1.233 | 0.2223 |
| *no burn vs. 1x* | 0.0268 | 0.187 | 0.8525 |
| *no burn vs. 2x* | -0.1848 | -1.287 | 0.2029 |
| *removal vs. reference* | -0.2499 | -2.122 | 0.038* |
| *removal vs. switch* | -0.2301 | -2.042 | 0.0456* |
| *removal vs. 1x* | -0.3833 | -3.499 | 0.0009*** |
| *removal vs. 2x* | -0.5949 | -5.432 | <0.0001*** |
| *reference vs. switch* | 0.0198 | 0.168 | 0.8668 |
| *reference vs. 1x* | -0.1333 | -1.166 | 0.2483 |
| *reference vs. 2x* | -0.345 | -3.016 | 0.0037** |
| *switch vs. 1x* | -0.1532 | -1.398 | 0.1671 |
| *switch vs. 2x* | -0.3648 | -3.331 | 0.0015** |
| *1x vs. 2x* | -0.2117 | -2.008 | 0.0492* |
| *p < 0.05, **p < 0.01, *** p < 0.0001 | | | |

**Section 6: Soil heating effects on plant communities**

**Table S10:** Contrasts for plant community composition PERMANOVA model.

| contrast | p-value |
| --- | --- |
| *no burn vs. removal* | 0.044* |
| *no burn vs. reference* | 0.044* |
| *no burn vs. switch* | 0.044* |
| *no burn vs. 1x* | 0.044* |
| *no burn vs. 2x* | 0.044* |
| *removal vs. reference* | 0.552 |
| *removal vs. switch* | 0.966 |
| *removal vs. 1x* | 0.966 |
| *removal vs. 2x* | 0.966 |
| *reference vs. switch* | 0.346 |
| *reference vs. 1x* | 0.333 |
| *reference vs. 2x* | 0.106 |
| *switch vs. 1x* | 0.966 |
| *switch vs. 2x* | 0.792 |
| *1x vs. 2x* | 0.966 |
| *p < 0.05, **p < 0.01, *** p < 0.0001 | |

**Table S11:** LMER model for soil heating and pine proximity effects on plant community species richness.

| model term | d.f. 1 | d.f. 2 | F-test | p-value |
| --- | --- | --- | --- | --- |
| *fuel load treatment* | 5 | 37.07 | 13.51 | <0.0001*** |
| *pines* | 1 | 16.14 | 0.269 | 0.6112 |
| *treatment:pines* | 5 | 30.07 | 0.815 | 0.5485 |
| random effect | variance | std. dev. |  |  |
| *patch within fire unit* | 6.2465 | 2.4993 |  |  |
| *fire unit* | 0.1254 | 0.3542 |  |  |
| *residual* | 8.0377 | 2.8351 |  |  |
| *p < 0.05, **p < 0.01, *** p < 0.0001 | | | | |

**Table S12:** Contrasts for plant community species richness LMER model.

| contrast | estimate | s.e. | d.f. | t-ratio | p-value |
| --- | --- | --- | --- | --- | --- |
| *no burn vs. removal* | 2.73 | 2.84 | 37.1 | 0.964 | 0.3412 |
| *no burn vs. reference* | -4.39 | 2.84 | 37.1 | -1.549 | 0.13 |
| *no burn vs. switch* | 5.23 | 2.84 | 37.1 | 1.846 | 0.0729 |
| *no burn vs. 1x* | 7.86 | 2.84 | 37.1 | 2.772 | 0.0087** |
| *no burn vs. 2x* | 10.48 | 2.84 | 37.1 | 3.697 | 0.0007** |
| *removal vs. reference* | -7.12 | 2 | 71.7 | -3.554 | 0.0007** |
| *removal vs. switch* | 2.5 | 2 | 71.7 | 1.247 | 0.2164 |
| *removal vs. 1x* | 5.12 | 2 | 71.7 | 2.556 | 0.0127* |
| *removal vs. 2x* | 7.75 | 2 | 71.7 | 3.866 | 0.0002** |
| *reference vs. switch* | 9.62 | 2 | 71.7 | 4.801 | <0.0001*** |
| *reference vs. 1x* | 12.25 | 2 | 71.7 | 6.111 | <0.0001*** |
| *reference vs. 2x* | 14.88 | 2 | 71.7 | 7.42 | <0.0001*** |
| *switch vs. 1x* | 2.62 | 2 | 71.7 | 1.309 | 0.1946 |
| *switch vs. 2x* | 5.25 | 2 | 71.7 | 2.619 | 0.0108* |
| *1x vs. 2x* | 2.62 | 2 | 71.7 | 1.309 | 0.1946 |
| *p < 0.05, **p < 0.01, *** p < 0.0001 | | | | | |

**Section 7: Soil heating effects on fungal communities**

**Table S13:** Pre vs. post fire contrasts for fungal community PERMANOVA model.

| Pine proximity | Fuel treat. | p-value |
| --- | --- | --- |
| near | removal | 0.0085** |
|  | reference | 0.0107* |
|  | switch | 0.00035*** |
|  | add 1x | 0.00035*** |
|  | add 2x | 0.00035*** |
| away | removal | 0.0171* |
|  | reference | 0.0118* |
|  | switch | 0.0025** |
|  | add 1x | 0.0039** |
|  | add 2x | 0.0074** |
| *p < 0.05, **p < 0.01, *** p < 0.0001 | | |

**Table S14:** Community type x sampling time contrasts for fungal community PERMANOVA model. Lower case letters denote significant differences (p<0.05) between fire severity groups within the same sampling time.

| Community type | Time | Fuel treatment | Significance group |
| --- | --- | --- | --- |
| litter | pre | removal | abc |
|  |  | reference | ab |
|  |  | switch | c |
|  |  | add 1x | abc |
|  |  | add 2x | abc |
|  | post | no burn | a |
|  |  | removal | b |
|  |  | reference | b |
|  |  | switch | bc |
|  |  | add 1x | c |
|  |  | add 2x | c |
| soil | pre | removal | ae |
|  |  | reference | bce |
|  |  | switch | bd |
|  |  | add 1x | ad |
|  |  | add 2x | abe |
|  | post | no burn | a |
|  |  | removal | bc |
|  |  | reference | ab |
|  |  | switch | bc |
|  |  | add 1x | cd |
|  |  | add 2x | d |

**Table S15:** LMER results for soil heating and pine proximity effects on fungal community diversity (Inverse Simpson Metric).

| model term | d.f. 1 | d.f. 2 | F-test | p-value |
| --- | --- | --- | --- | --- |
| fuel load treatment | 5 | 83.06 | 6.988 | <0.0001*** |
| pine proximity | 1 | 7.33 | 2.477 | 0.1576 |
| community type | 1 | 141.39 | 8.004 | 0.0053** |
| fire x pine | 5 | 83.42 | 1.45 | 0.2151 |
| fire x comm. type | 5 | 143.55 | 2.296 | 0.0484* |
| pine x comm. type | 1 | 141.47 | 6.888 | 0.0096** |
| fire x pine x comm. | 5 | 143.81 | 4.692 | 0.0005*** |
| random effect | variance | std. dev. |  |  |
| *patch within fire unit* | 37.47 | 6.121 |  |  |
| *fire unit* | 25.97 | 5.096 |  |  |
| *Residual* | 1061.39 | 32.579 |  |  |
| *p < 0.05, **p < 0.01, *** p < 0.0001 | | | | |

**Table S16:** Contrasts for fungal diversity LMER model.

| Community type | contrast | estimate | s.e. | d.f. | t-ratio | p-value |
| --- | --- | --- | --- | --- | --- | --- |
|  | *litter vs. soil* | -174.36 | 61.6 | 141 | -2.829 | 0.0053 |
| litter | *near vs. away* | -140.66 | 46.7 | 17 | -3.014 | 0.0078** |
|  | *no burn vs. removal* | 65.85 | 24.1 | 125 | 2.727 | 0.0073** |
|  | *no burn vs. reference* | 98.98 | 23 | 146 | 4.3 | <0.0001*** |
|  | *no burn vs. switch* | 95.9 | 25.4 | 130 | 3.777 | 0.0002*** |
|  | *no burn vs. 1x* | 133.64 | 23.7 | 124 | 5.631 | <0.0001*** |
|  | *no burn vs. 2x* | 122.08 | 23.7 | 124 | 5.144 | <0.0001*** |
|  | *removal vs. reference* | 51.91 | 23.5 | 139 | 2.212 | 0.0286* |
|  | *removal vs. switch* | 30.06 | 25.1 | 139 | 1.196 | 0.2337 |
|  | *removal vs. 1x* | 67.79 | 23.5 | 139 | 2.889 | 0.0045** |
|  | *removal vs. 2x* | 56.23 | 23.5 | 139 | 2.397 | 0.0179* |
|  | *reference vs. switch* | -21.85 | 24.7 | 139 | -0.883 | 0.3786 |
|  | *reference vs. 1x* | 15.88 | 23 | 138 | 0.689 | 0.4918 |
|  | *reference vs. 2x* | 4.33 | 23 | 138 | 0.188 | 0.8513 |
|  | *switch vs. 1x* | 37.73 | 24.7 | 139 | 1.525 | 0.1295 |
|  | *switch vs. 2x* | 26.18 | 24.7 | 139 | 1.058 | 0.2918 |
|  | *1x vs. 2x* | -11.55 | 23 | 138 | -0.501 | 0.6168 |
| soil | *near vs. away* | 21.39 | 51 | 22.5 | 0.42 | 0.6788 |
|  | *no burn vs. removal* | -29.66 | 29.9 | 132 | -0.992 | 0.3229 |
|  | *no burn vs. reference* | -55.79 | 27.4 | 147 | -2.033 | 0.0439* |
|  | *no burn vs. switch* | 23.82 | 29.9 | 132 | 0.798 | 0.4266 |
|  | *no burn vs. 1x* | 38.22 | 29.5 | 131 | 1.293 | 0.1981 |
|  | *no burn vs. 2x* | 42.65 | 29.9 | 132 | 1.426 | 0.1561 |
|  | *removal vs. reference* | 23.05 | 26 | 139 | 0.886 | 0.3771 |
|  | *removal vs. switch* | 53.48 | 23.9 | 139 | 2.239 | 0.0267* |
|  | *removal vs. 1x* | 67.88 | 23.5 | 139 | 2.893 | 0.0044** |
|  | *removal vs. 2x* | 72.31 | 23.9 | 139 | 3.028 | 0.0029** |
|  | *reference vs. switch* | 30.44 | 26 | 139 | 1.17 | 0.2439 |
|  | *reference vs. 1x* | 44.84 | 25.6 | 139 | 1.748 | 0.0826 |
|  | *reference vs. 2x* | 49.27 | 26 | 139 | 1.893 | 0.0604 |
|  | *switch vs. 1x* | 14.4 | 23.5 | 139 | 0.614 | 0.5404 |
|  | *switch vs. 2x* | 18.83 | 23.9 | 139 | 0.788 | 0.4321 |
|  | *1x vs. 2x* | 4.43 | 23.5 | 139 | 0.189 | 0.8505 |
| *p < 0.05, **p < 0.01, *** p < 0.0001 | | | | | |  |

**Table S17:** Contrasts for fungal community heterogeneity ANOVA model.

| contrast | estimate | lower c.i. | upper c.i. | p-value |
| --- | --- | --- | --- | --- |
| *no burn vs. removal* | -7.59731 | -16.2757 | 1.081116 | 0.1238382 |
| *no burn vs. reference* | -8.914 | -17.6302 | -0.19783 | 0.041713* |
| *no burn vs. switch* | -11.5673 | -20.3228 | -2.81178 | 0.0025421** |
| *no burn vs. 1x* | -14.9751 | -23.6535 | -6.29667 | <0.0001*** |
| *no burn vs. 2x* | -16.8635 | -25.5796 | -8.1473 | <0.0001*** |
| *removal vs. reference* | -1.31669 | -8.96081 | 6.327419 | 0.9963246 |
| *removal vs. switch* | -3.96996 | -11.6589 | 3.718959 | 0.6753554 |
| *removal vs. 1x* | -7.37779 | -14.9788 | 0.223259 | 0.0627635 |
| *removal vs. 2x* | -9.26616 | -16.9103 | -1.62205 | 0.0076884** |
| *reference vs. switch* | -2.65327 | -10.3848 | 5.078229 | 0.9221452 |
| *reference vs. 1x* | -6.0611 | -13.7052 | 1.583018 | 0.207234 |
| *reference vs. 2x* | -7.94947 | -15.6364 | -0.26253 | 0.0379987* |
| *switch vs. 1x* | -3.40782 | -11.0967 | 4.281098 | 0.799396 |
| *switch vs. 2x* | -5.2962 | -13.0277 | 2.435301 | 0.3639608 |
| *1x vs. 2x* | -1.88837 | -9.53249 | 5.755738 | 0.9806621 |
| *p < 0.05, **p < 0.01, *** p < 0.0001 | | | | |

**Description of indicator species**

Prior to fire, taxa like *Pyrenochaetopsis leptospora* (plant pathogen; tables S22-31), *Cenococcum geophilum* (ectomycorrhizal), *Parateratosphaerie altensteinii* (plant pathogen), *Lophodermium conigenum* (pine pathogen), *Phialophora livistonae* (foliage associated), and members of *Catenulifera* (mycoparasites) were higher in abundance relative to post-fire communities. Following fire however, members of *Talaromyces* (thermophilic), *Aureobasidium* (black yeast), *Papiliotrema perniciosus* (oxidation resistant), and *Curvibasidium cygneicollum* (degrades combustion products) increased in abundance. While there were similar trends in indicator taxa across fire severity treatments, the number of indicator taxa decreased in lower fire severity treatments, and were lower in soil relative to litter fungal communities.

**Table S18:** Pre vs. post fire indicator taxa for 2X pine needle edition treatments.

| Time | ASV | Natural history | citation |
| --- | --- | --- | --- |
|  |  | litter indicator species |  |
| pre | Pyrenochaetopsis leptospora | Many *pyrenochaetopsis* are plant pathogens, saprotrophs, and endophytes | (da Silva *et al.*, 2019) |
|  | Cenococcum geophilum | Common ectomychorrizal fungus associated with pine species | (Trappe, 1962) |
|  | Catenulifera sp. |  |  |
|  | Parateratosphaeria altensteinii | Causes leaf spots, likely plant pathogen | (Crous, 2017) |
|  | Lophiostomataceae sp. | Likely saprotroph | (Cannon & Kirk, 2007) |
|  | Pleosporales sp. |  |  |
|  | Chaetothyriales sp. |  |  |
|  | Unclassifed fungus |  |  |
|  | Unclassifed fungus |  |  |
|  | Periconia byssoides | Likely saprotroph, found in mowed grassland sites | (Hopkins *et al.*, 2021) |
|  | Ascomycota sp. |  |  |
| post | Curvibasidium cygneicollum | some can degrade benzo(a)anthrace, a combustion byproduct | (MacGillivray & Shiaris, 1993) |
|  | Didymellaceae sp. | Likely plant pathogen |  |
|  | Talaromyces sp. | *Talaromyces* range from mesophilic to strongly thermophilic | (Stolk & Samson, 1972) |
|  | Sordariaceae sp. | Contains dark, mold forming species. Some are coprophilous |  |
|  | Aureobasidium sp. | Genus is composed of black, yeast like fungi | (Humphries *et al.*, 2017) |
|  | Didymellaceae sp. | Likely plant pathogen |  |
|  | Cladosporium sp. | mold, saprotroph | (Deshmukh, S.K. & Rai, 2005) |
|  | Aureobasidium namibiae | Black, yeast like fungus | (Humphries *et al.*, 2017) |
|  | Papiliotrema perniciosus | Used as biocontrol agent of yeasts, resistant to oxidative stress | (Palmieri *et al.*, 2021) |
|  | Filobasidium magnum |  |  |
|  |  | soil indicator species |  |
| pre | none |  |  |
| post | Didymellaceae sp. | Likely plant pathogen |  |
|  | Talaromyces sp. | *Talaromyces* range from mesophilic to strongly thermophilic | (Stolk & Samson, 1972) |
|  | Didymellaceae sp. | Likely plant pathogen |  |
|  | Alternaria sp. | Likely plant pathogen | (Cannon & Kirk, 2007) |

**Table S19:** ALDEx2 results for 2X fuel load treatments. Diff.btw is the median difference between groups on a log base 2 scale. Diff.win is the largest median variation within group. Effect is the effect size of diff.btw/diff.win and describes whether inter- vs. intra-group variance is larger. Overlap describes confusion in assigning an observation to either group. Wi.ep is the expected value of the Wilcoxon test p-value. Wi.eBH is the expected value of the Benjamini-Hochberg corrected p-value.

| Time | ASV | wi.ep | wi.eBH | diff.btw | diff.win | effect | overlap |
| --- | --- | --- | --- | --- | --- | --- | --- |
| litter indicator taxa | | | | | | | |
| pre | Pyrenochaetopsis leptospora | 1.88E-03 | 0.06803 | 6.359057 | 4.959852 | 1.175764 | 0.071652 |
|  | Cenococcum geophilum | 2.98E-03 | 0.086585 | 6.229896 | 4.008624 | 1.553743 | 0.071876 |
|  | Catenulifera sp. | 8.78E-04 | 0.047337 | 8.165971 | 4.581382 | 1.617242 | 0.040501 |
|  | Parateratosphaeria altensteinii | 9.83E-05 | 0.017384 | 7.99181 | 2.864345 | 2.602403 | 0.000219 |
|  | Lophiostomataceae sp. | 1.60E-03 | 0.062379 | 6.544453 | 3.295341 | 1.799367 | 0.059377 |
|  | Pleosporales sp. | 2.64E-04 | 0.024974 | 7.012612 | 3.642011 | 1.730478 | 0.015583 |
|  | Chaetothyriales sp. | 1.54E-04 | 0.019212 | 6.012135 | 3.467107 | 1.576676 | 0.012469 |
|  | Unclassifed fungus | 2.12E-03 | 0.072851 | 7.435739 | 3.165601 | 1.873078 | 0.062502 |
|  | Unclassifed fungus | 9.83E-05 | 0.017384 | 8.108863 | 2.602002 | 3.116951 | 0.000219 |
|  | Periconia byssoides | 2.78E-04 | 0.024535 | 5.945154 | 2.710105 | 2.230244 | 0.018755 |
|  | Ascomycota sp. | 2.58E-04 | 0.022776 | 6.02526 | 2.988247 | 1.933921 | 0.009356 |
| post | Curvibasidium cygneicollum | 0.00068 | 0.037269 | -11.5288 | 3.964611 | -2.77298 | 0.031253 |
|  | Didymellaceae sp. | 0.003315 | 0.085083 | -11.7301 | 3.895929 | -2.62397 | 0.065422 |
|  | Talaromyces sp. | 0.006582 | 0.117557 | -9.66043 | 4.488323 | -1.67763 | 0.080998 |
|  | Curvibasidium cygneicollum | 0.008171 | 0.137833 | -9.95657 | 4.853169 | -1.49146 | 0.109035 |
|  | Sordariaceae sp. | 0.011721 | 0.159513 | -10.2989 | 4.749561 | -1.72974 | 0.102805 |
|  | Aureobasidium sp. | 0.006455 | 0.1174 | -9.65099 | 4.154871 | -1.8568 | 0.090344 |
|  | Didymellaceae sp. | 0.000653 | 0.036032 | -9.74842 | 3.099484 | -2.97889 | 0.034271 |
|  | Cladosporium sp. | 0.032046 | 0.260533 | -8.78396 | 6.507438 | -1.07596 | 0.165625 |
|  | Aureobasidium namibiae | 0.005393 | 0.10708 | -8.75551 | 3.950584 | -1.8527 | 0.080998 |
|  | Papiliotrema perniciosus | 0.000752 | 0.038627 | -8.52636 | 3.544375 | -2.30491 | 0.031156 |
|  | Filobasidium magnum | 0.002374 | 0.068277 | -6.57153 | 3.624875 | -1.80084 | 0.052961 |
| soil indicator taxa | | | | | | | |
| pre | none | - | - | - | - | - | - |
| post | Didymellaceae sp. | 4.08E-06 | 0.002244 | -10.3499 | 3.918812 | -2.52515 | 0.000137 |
|  | Talaromyces sp. | 1.82E-04 | 0.040888 | -3.66237 | 1.61054 | -1.94733 | 0.060547 |
|  | Didymellaceae sp. | 4.08E-06 | 0.002244 | -10.0903 | 3.290039 | -3.04364 | 0.000137 |
|  | Alternaria sp. | 1.91E-03 | 0.1233 | -7.35263 | 4.519508 | -1.49283 | 0.082032 |

**Table S20:** Pre vs. post fire indicator taxa for 1X pine needle edition treatments.

| Time | ASV | Natural history | citation |
| --- | --- | --- | --- |
| litter indicator taxa | | | |
| pre | Cenococcum geophilum | Common ectomychorrizal fungus associated with pine species | (Trappe, 1962) |
|  | Catenulifera sp. | Genus contains saprotrophs and mycoparasites | (Bogale *et al.*, 2010) |
|  | Seiridium sp. | Can cause Seiridium cankers in trees | (Della Rocca *et al.*, 2011) |
|  | Chaetothyriales sp. |  |  |
|  | Unclassified fungus |  |  |
| post | none |  |  |
| soil indicator taxa | | | |
| pre | none |  |  |
| post | Didymellaceae sp. | Likely plant pathogen |  |
|  | Curvibasidium cygneicollum | some Rhodotorula can degrade benzo(a)anthrace, a combustion byproduct | (MacGillivray & Shiaris, 1993) |
|  | Didymellaceae sp. | Likely plant pathogen |  |

**Table S21:** ALDEx2 results for 1X fuel load treatments. Diff.btw is the median difference between groups on a log base 2 scale. Diff.win is the largest median variation within group. Effect is the effect size of diff.btw/diff.win and describes whether inter- vs. intra-group variance is larger. Overlap describes confusion in assigning an observation to either group. Wi.ep is the expected value of the Wilcoxon test p-value. Wi.eBH is the expected value of the Benjamini-Hochberg corrected p-value.

| Time | ASV | wi.ep | wi.eBH | diff.btw | diff.win | effect | overlap |
| --- | --- | --- | --- | --- | --- | --- | --- |
| litter indicator taxa | |  |  |  |  |  |  |
| pre | Cenococcum geophilum | 0.006872 | 0.2215 | 6.935086 | 4.477355 | 1.22027 | 0.093752 |
|  | Catenulifera sp. | 0.004737 | 0.191299 | 4.964978 | 4.988476 | 1.01765 | 0.062259 |
|  | Seiridium sp. | 0.004921 | 0.197068 | 8.751128 | 4.509754 | 1.898776 | 0.062502 |
|  | Chaetothyriales sp. | 0.023739 | 0.367716 | 6.136938 | 4.177166 | 1.208874 | 0.116733 |
|  | Unclassified fungus | 0.000413 | 0.052815 | 7.853209 | 3.386395 | 2.204303 | 0.000274 |
| post | none | - | - | - | - | - | - |
| soil indicator taxa | |  |  |  |  |  |  |
| pre | none | - | - | - | - | - | - |
| post | Didymellaceae sp. | 5.06E-06 | 0.00619 | -10.1627 | 3.884179 | -2.44488 | 0.019066 |
|  | Curvibasidium cygneicollum | 1.30E-03 | 0.161034 | -7.47481 | 4.335699 | -1.48406 | 0.097223 |
|  | Didymellaceae sp. | 1.53E-04 | 0.04548 | -9.3391 | 3.847212 | -2.17783 | 0.052084 |

**Table S22:** Pre vs. post fire indicator taxa for switch fuel load treatments.

| Time | ASV | Natural history | citation |
| --- | --- | --- | --- |
| litter indicator taxa | | | |
| pre | Strelitziana syzygii | Member of Trichomeriaceae, sooty moulds, leaf and insect exudate associated | (Chomnunti *et al.*, 2012) |
|  | Phialophora livistonae | Dark septate endophyte, previously found in non-burned pine savannas and prairies | Crous and Summerell 2012 (Hopkins *et al.*, 2021) |
|  | Unclassified fungus |  |  |
|  | Lophodermium conigenum | Forms black spots on pine needles, associated with *P. sylvestris* | (Minter *et al.*, 1978) |
|  | Pleosporales sp. |  |  |
|  | Periconia byssoides | Likely saprotroph, found in mowed grassland sites | (Hopkins *et al*., 2021) |
|  | Chaetothyriales sp. |  |  |
| post | Didymellaceae sp. | Likely plant pathogen |  |
|  | Curvibasidium cygneicollum | some Rhodotorula can degrade benzo(a)anthrace, a combustion byproduct | (MacGillivray & Shiaris, 1993) |
|  | Tremella sp. | All tremella are mycoparasites | (Chen, 1998) |
|  | Tremella sp. | All tremella are mycoparasites | (Chen, 1998) |
| soil indicator taxa | | | |
| pre | none | - | - |
| post | none | - | - |

**Table S23:** ALDEx2 results for switch fuel load treatments. Diff.btw is the median difference between groups on a log base 2 scale. Diff.win is the largest median variation within group. Effect is the effect size of diff.btw/diff.win and describes whether inter- vs. intra-group variance is larger. Overlap describes confusion in assigning an observation to either group. Wi.ep is the expected value of the Wilcoxon test p-value. Wi.eBH is the expected value of the Benjamini-Hochberg corrected p-value.

| Time | ASV | wi.ep | wi.eBH | diff.btw | diff.win | effect | overlap |
| --- | --- | --- | --- | --- | --- | --- | --- |
| litter indicator taxa | | | | | | | |
| pre | Strelitziana syzygii | 2.82E-05 | 0.004955 | 7.336293 | 3.376882 | 2.068496 | 0.002255 |
|  | Phialophora livistonae | 2.58E-05 | 0.004702 | 8.284053 | 2.816468 | 2.777108 | 0.000157 |
|  | unidentified | 5.38E-04 | 0.043069 | 7.050129 | 3.913378 | 1.56523 | 0.055804 |
|  | Lophodermium conigenum | 2.86E-03 | 0.104766 | 9.674996 | 3.91428 | 2.132864 | 0.089087 |
|  | Pleosporales sp. | 2.58E-05 | 0.004702 | 8.116353 | 2.683514 | 3.315985 | 0.000157 |
|  | Periconia byssoides | 1.01E-04 | 0.012182 | 6.793831 | 3.320508 | 1.84332 | 0.024556 |
|  | Chaetothyriales sp. | 2.58E-05 | 0.004702 | 7.956919 | 3.070992 | 2.475701 | 0.000157 |
|  | Periconia byssoides | 6.99E-05 | 0.009053 | 7.159063 | 3.3595 | 2.045075 | 0.008914 |
| post | Didymellaceae sp. | 2.58E-05 | 0.004702 | -10.0058 | 4.908429 | -1.83666 | 0.000157 |
|  | Curvibasidium cygneicollum | 2.58E-05 | 0.004702 | -10.8735 | 3.065684 | -3.5676 | 0.000157 |
|  | Tremella sp. | 1.56E-03 | 0.084428 | -2.68402 | 1.649729 | -1.52325 | 0.064589 |
|  | Tremella sp. | 2.03E-03 | 0.080736 | -8.63128 | 3.607534 | -1.957 | 0.091518 |
| soil indicator taxa | | | | | | | |
| pre | none | - | - | - | - | - | - |
| post | none | - | - | - | - | - | - |

**Table S24:** Pre vs. post fire indicator taxa for reference fuel load treatments.

| Time | ASV | Natural history | citation |
| --- | --- | --- | --- |
|  |  | litter indicator taxa |  |
| pre | Strelitziana syzygii | Member of Trichomeriaceae, sooty moulds, leaf and insect exudate associated | (Chomnunti *et al.*, 2012) |
|  | Phialophora livistonae | Dark septate endophyte, previously found in non-burned pine savannas and prairies | (Crous & Summerell, 2012) (Hopkins *et al.*, 2021) |
| post | Didymellaceae sp. | Likely plant pathogen |  |
|  | Curvibasidium cygneicollum | some Rhodotorula can degrade benzo(a)anthrace, a combustion byproduct | (MacGillivray & Shiaris, 1993) |
|  | Didymellaceae sp. | Likely plant pathogen |  |
|  |  | soil indicator taxa |  |
| pre | none | - | - |
| post | none | - | - |

**Table S25:** ALDEx2 results for reference fuel load treatments. Diff.btw is the median difference between groups on a log base 2 scale. Diff.win is the largest median variation within group. Effect is the effect size of diff.btw/diff.win and describes whether inter- vs. intra-group variance is larger. Overlap describes confusion in assigning an observation to either group. Wi.ep is the expected value of the Wilcoxon test p-value. Wi.eBH is the expected value of the Benjamini-Hochberg corrected p-value.

| Time | ASV | wi.ep | wi.eBH | diff.btw | diff.win | effect | overlap |
| --- | --- | --- | --- | --- | --- | --- | --- |
| pre | Strelitziana syzygii | 0.000123 | 0.032036 | 8.612905 | 3.949959 | 1.954135 | 0.046876 |
|  | Phialophora livistonae | 0.000627 | 0.072157 | 7.697856 | 3.894396 | 1.70013 | 0.095517 |
|  | Phialophora livistonae | 0.000479 | 0.048026 | 6.326567 | 3.283282 | 1.778004 | 0.062379 |
| post | Didymellaceae sp. | 6.73E-06 | 0.004902 | -10.3938 | 4.687893 | -1.93879 | 0.007802 |
|  | Curvibasidium cygneicollum | 5.74E-06 | 0.004902 | -10.1777 | 4.305451 | -2.21643 | 0.005866 |
|  | Didymellaceae sp. | 2.10E-05 | 0.009411 | -9.69268 | 4.707769 | -1.79586 | 0.029241 |
| pre | none | - | - | - | - | - | - |
| post | none | - | - | - | - | - | - |

**Table S26:** Pre vs. post fire indicator taxa for removal fuel load treatments.

| Time | ASV | Natural history | citation |
| --- | --- | --- | --- |
|  |  | litter indicator taxa |  |
| pre | Phialophora livistonae | Dark septate endophyte, found in non-burned pine savannas and prairies | (Crous & Summerell 2012) (Hopkins *et al.,* 2021) |
|  | Pleosporales sp. |  |  |
|  | Periconia byssoides | Likely saprotroph, found in mowed grassland sites | (Hopkins *et al.,* 2021) |
|  | Chaetothyriales sp. |  |  |
|  | Strelitziana syzygii | Member of Trichomeriaceae, sooty moulds, leaf and insect associated | (Chomnunti *et al.*, 2012) |
|  | Lophiostomataceae sp. | Likely saprotroph | (Cannon and Kirk, 2007) |
|  | Unclassified fungus |  |  |
|  | Xenomycosphaerella elongata | Likely a plant pathogen | (Quaedvlieg, W. *et al.*, 2014) |
|  | Colletotrichum gloeosporioides | Plant pathogen associated with fruits, causes Anthracnose | (Siddiqui & Ali, 2014) |
| post | Curvibasidium cygneicollum | some Rhodotorula can degrade benzo(a)anthrace, a combustion byproduct | (MacGillivray & Shiaris, 1993) |
|  | Didymellaceae sp. | Likely plant pathogen |  |
|  | Aureobasidium sp. | Genus is composed of black, yeast like fungi | (Humphries *et al.*, 2017) |
|  |  | soil indicator taxa |  |
| pre | none | - | - |
| post | none | - | - |

**Table S27:** ALDEx2 results for removal fuel load treatments. Diff.btw is the median difference between groups on a log base 2 scale. Diff.win is the largest median variation within group. Effect is the effect size of diff.btw/diff.win and describes whether inter- vs. intra-group variance is larger. Overlap describes confusion in assigning an observation to either group. Wi.ep is the expected value of the Wilcoxon test p-value. Wi.eBH is the expected value of the Benjamini-Hochberg corrected p-value.

| Time | ASV | wi.ep | wi.eBH | diff.btw | diff.win | effect | overlap |
| --- | --- | --- | --- | --- | --- | --- | --- |
| litter indicator taxa | | | | | | | |
| pre | Phialophora livistonae | 5.25E-04 | 0.045435 | 5.745041 | 4.30837 | 1.180072 | 0.066816 |
|  | Pleosporales sp. | 5.52E-05 | 0.011377 | 7.375202 | 4.048003 | 1.638602 | 0.024556 |
|  | Periconia byssoides | 4.09E-05 | 0.009847 | 6.437853 | 4.151345 | 1.473087 | 0.022324 |
|  | Chaetothyriales sp. | 9.37E-06 | 0.005032 | 7.366456 | 3.201226 | 2.18814 | 0.000157 |
|  | Strelitziana syzygii | 9.76E-04 | 0.053129 | 8.19344 | 3.582977 | 1.926757 | 0.066965 |
|  | Lophiostomataceae sp. | 1.39E-03 | 0.066725 | 7.912839 | 3.942977 | 1.861513 | 0.073497 |
|  | Unclassified fungus | 6.28E-05 | 0.012691 | 6.611687 | 3.778493 | 1.676283 | 0.026728 |
|  | Xenomycosphaerella elongata | 1.95E-03 | 0.081679 | 7.516643 | 4.454421 | 1.471855 | 0.075893 |
|  | Colletotrichum gloeosporioides | 1.09E-03 | 0.056316 | 6.706212 | 3.758367 | 1.690613 | 0.062501 |
| post | Curvibasidium cygneicollum | 8.22E-06 | 0.004966 | -10.5672 | 3.893048 | -2.7136 | 0.000157 |
|  | Didymellaceae sp. | 7.06E-04 | 0.049194 | -9.58258 | 4.353522 | -1.99952 | 0.060134 |
|  | Curvibasidium cygneicollum | 4.05E-03 | 0.135248 | -9.31717 | 4.509171 | -1.59503 | 0.111608 |
|  | Aureobasidium sp. | 6.96E-03 | 0.166133 | -8.30107 | 5.459777 | -1.31794 | 0.124722 |
| soil indicator taxa | | | | | | | |
| pre | none | - | - | - | - | - | - |
| post | none | - | - | - | - | - | - |

**Section 8: Soil heating effects on microbial decomposition -**

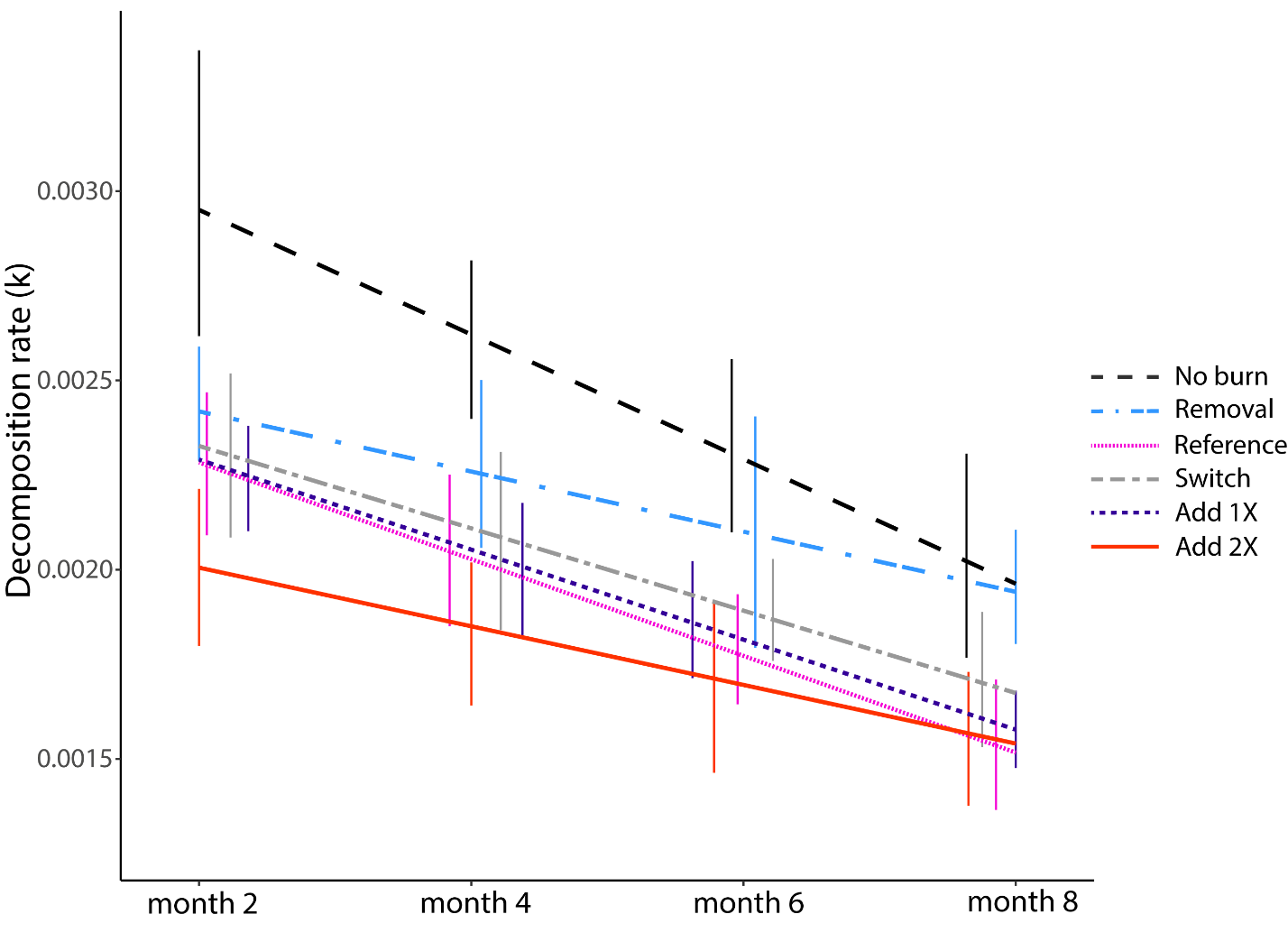

**Figure S28:** Fuel manipulation treatment effects on microbial decomposition rate (k). Error bars represent +/- one S.E.. Trend lines describe k-rates across the eight months following 2017 prescribed burns. Microbial decomposition was slower in burned relative to no burn plots during the study period. Early after fire, treatments with larger fuel loads displayed the lowest k-rates.

**Table S29:** Contrasts for microbial decomposition rate LMER models.

|  | 2 month k-rate | | |  | 4 month k-rate | | |
| --- | --- | --- | --- | --- | --- | --- | --- |
| contrast | estimate | t-ratio | p-value |  | estimate | t-ratio | p-value |
| *away vs. near* | -0.04553 | -0.069 | 0.9469 |  | 2.65E-04 | 0.247 | 0.8058 |
| *fire vs. no fire* | -2.65829 | -2.26 | 0.0367* |  | -4.05E-03 | -1.38 | 0.1721 |
| *no burn vs. removal* | 0.52769 | 2.061 | 0.0504* |  | 6.68E-05 | 0.098 | 0.9224 |
| *no burn vs. reference* | 0.43143 | 1.627 | 0.1155 |  | 1.21E-03 | 1.666 | 0.1004 |
| *no burn vs. switch* | 0.41954 | 1.641 | 0.114 |  | 6.45E-04 | 0.944 | 0.3486 |
| *no burn vs. 1x* | 0.52497 | 2.067 | 0.0501* |  | 9.48E-04 | 1.403 | 0.1651 |
| *no burn vs. 2x* | 0.75467 | 2.909 | 0.0076* |  | 1.18E-03 | 1.707 | 0.0924. |
| *Removal vs. reference* | -0.09625 | -0.549 | 0.585 |  | 1.14E-03 | 1.874 | 0.0653. |
| *Removal vs. switch* | -0.10814 | -0.672 | 0.5043 |  | 5.78E-04 | 1.035 | 0.3045 |
| *Removal vs. 1x* | -0.00272 | -0.017 | 0.9863 |  | 8.81E-04 | 1.604 | 0.1133 |
| *Removal vs. 2x* | 0.22698 | 1.369 | 0.1763 |  | 1.12E-03 | 1.956 | 0.0546* |
| *reference vs. switch* | -0.01189 | -0.068 | 0.9462 |  | -5.61E-04 | -0.923 | 0.3593 |
| *reference vs. 1x* | 0.09354 | 0.541 | 0.5906 |  | -2.58E-04 | -0.431 | 0.6681 |
| *reference vs. 2x* | 0.32323 | 1.799 | 0.0772. |  | -2.24E-05 | -0.036 | 0.9712 |
| *switch vs. 1x* | 0.10542 | 0.667 | 0.5072 |  | 3.03E-04 | 0.552 | 0.5829 |
| *switch vs. 2x* | 0.33512 | 2.014 | 0.0487* |  | 5.39E-04 | 0.944 | 0.3487 |
| *1x vs. 2x* | 0.2297 | 1.41 | 0.1641 |  | 2.36E-04 | 0.42 | 0.6761 |
|  | 6 month k-rate | | |  | 8 month k-rate | | |
| *away vs. near* | 1.67E-03 | 1.52 | 0.1332 |  | -1.54E-03 | -0.719 | 0.4823 |
| *fire vs. no fire* | -2.64E-03 | -0.876 | 0.384 |  | 6.25E-03 | 1.257 | 0.2131 |
| *no burn vs. removal* | -2.84E-04 | -0.406 | 0.6864 |  | -6.46E-04 | -0.568 | 0.5718 |
| *no burn vs. reference* | 8.44E-04 | 1.136 | 0.2599 |  | -1.41E-03 | -1.174 | 0.2437 |
| *no burn vs. switch* | 8.90E-04 | 1.27 | 0.2086 |  | -6.23E-04 | -0.548 | 0.5851 |
| *no burn vs. 1x* | 2.00E-04 | 0.288 | 0.7741 |  | -2.08E-03 | -1.844 | 0.0687. |
| *no burn vs. 2x* | 9.89E-04 | 1.39 | 0.1692 |  | -1.50E-03 | -1.294 | 0.1992 |
| *Removal vs. reference* | 1.13E-03 | 1.809 | 0.075. |  | -7.64E-04 | -0.796 | 0.4285 |
| *Removal vs. switch* | 1.17E-03 | 2.049 | 0.0444* |  | 2.23E-05 | 0.025 | 0.9799 |
| *Removal vs. 1x* | 4.84E-04 | 0.859 | 0.3935 |  | -1.43E-03 | -1.652 | 0.1026 |
| *Removal vs. 2x* | 1.27E-03 | 2.173 | 0.0333* |  | -8.52E-04 | -0.939 | 0.3504 |
| *reference vs. switch* | 4.59E-05 | 0.074 | 0.9416 |  | 7.86E-04 | 0.818 | 0.4156 |
| *reference vs. 1x* | -6.45E-04 | -1.048 | 0.2985 |  | -6.65E-04 | -0.702 | 0.4845 |
| *reference vs. 2x* | 1.44E-04 | 0.227 | 0.821 |  | -8.75E-05 | -0.089 | 0.9293 |
| *switch vs. 1x* | -6.91E-04 | -1.225 | 0.225 |  | -1.45E-03 | -1.677 | 0.0974. |
| *switch vs. 2x* | 9.85E-05 | 0.168 | 0.867 |  | -8.74E-04 | -0.962 | 0.339 |
| *1x vs. 2x* | 7.89E-04 | 1.368 | 0.1758 |  | 5.78E-04 | 0.649 | 0.5184 |

. = p<0.1; * = p<0.05; ** = p<0.0001

**Section 9: Structural equation model results**

**Table S30:** Results from final structural equation model for soil heating effects on microbial decomposition.

| Dependent variable | Independent variable | Estimate | S.E. | z-value | p-value | Std. est. | R^2^ |
| --- | --- | --- | --- | --- | --- | --- | --- |
| soil heating (latent) | max. surf. temp. change | 1 |  |  |  | 0.714 | 0.51 |
|  | surf. dur. >60C | 0.213 | 0.192 | 1.109 | 0.268 | 0.14 | 0.02 |
|  | max. soil temp. change | 0.622 | 0.189 | 3.298 | 0.001* | 0.416 | 0.173 |
| soil heating | fuel load treatment | 0.422 | 0.057 | 7.448 | 0** | 0.889 | 0.956 |
|  | pine proximity | 0.508 | 0.138 | 3.681 | 0** | 0.363 |  |
| fine fuel combustion | soil heating | 0.966 | 0.204 | 4.738 | 0** | 0.662 | 0.38 |
|  | pine proximity | -0.798 | 0.257 | -3.105 | 0.002* | -0.391 |  |
| litter fungi PCA axis 1 | soil heating | -0.235 | 0.129 | -1.829 | 0.067. | -0.191 | 0.13 |
|  | plant comm. PCoA 1 | -0.165 | 0.065 | -2.539 | 0.011* | -0.201 |  |
|  | soil pH | -0.11 | 0.068 | -1.611 | 0.107 | -0.131 |  |
|  | plant comm. PCoA 2 | 0.042 | 0.062 | 0.685 | 0.494 | 0.048 |  |
| litter fungi PCA axis 2 | plant comm. PCoA 1 | -0.187 | 0.075 | -2.487 | 0.013* | -0.215 | 0.076 |
|  | plant species richness | 0.207 | 0.108 | 1.915 | 0.055* | 0.215 |  |
|  | C:N ratio | 0.076 | 0.087 | 0.875 | 0.382 | 0.085 |  |
|  | NH4+ | 0.126 | 0.073 | 1.737 | 0.082. | 0.14 |  |
|  | NO3- | 0.143 | 0.084 | 1.693 | 0.091. | 0.124 |  |
| litter fungal diversity | soil heating | -0.446 | 0.101 | -4.43 | 0** | -0.367 | 0.161 |
|  | plant species richness | -0.228 | 0.072 | -3.172 | 0.002* | -0.253 |  |
|  | C:N ratio | -0.11 | 0.059 | -1.844 | 0.065. | -0.13 |  |
| soil fungi PCA 1 | pine proximity | -0.797 | 0.179 | -4.442 | 0** | -0.404 | 0.42 |
|  | plant comm. PCoA 1 | 0.366 | 0.076 | 4.839 | 0** | 0.388 |  |
|  | C:N ratio | -0.189 | 0.08 | -2.361 | 0.018* | -0.194 |  |
|  | soil pH | -0.499 | 0.088 | -5.648 | 0** | -0.518 |  |
|  | NO3- | -0.216 | 0.097 | -2.212 | 0.027* | -0.173 |  |
| soil fungi PCA 2 | soil heating | -0.417 | 0.136 | -3.067 | 0.002* | -0.297 | 0.406 |
|  | plant comm. PCoA 2 | -0.352 | 0.086 | -4.11 | 0** | -0.348 |  |
|  | C:N ratio | -0.137 | 0.082 | -1.661 | 0.097. | -0.141 |  |
|  | NO3- | 0.431 | 0.1 | 4.295 | 0** | 0.348 |  |
| soil fungal diversity | pine proximity | -0.466 | 0.196 | -2.382 | 0.017* | -0.244 | 0.215 |
|  | plant comm. PCoA 2 | 0.095 | 0.085 | 1.117 | 0.264 | 0.096 |  |
|  | inorg. phosp. | 0.107 | 0.082 | 1.316 | 0.188 | 0.115 |  |
|  | soil pH | -0.419 | 0.097 | -4.336 | 0** | -0.45 |  |
|  | NH4+ | -0.13 | 0.079 | -1.655 | 0.098. | -0.138 |  |
| plant comm. PCoA 1 | inorg. phosp. | 0.651 | 0.083 | 7.892 | 0** | 0.637 | 0.577 |
|  | NH4+ | -0.325 | 0.083 | -3.899 | 0** | -0.315 |  |
|  | NO3- | 0.354 | 0.107 | 3.318 | 0.001** | 0.268 |  |
| plant species richness | soil heating | -0.592 | 0.196 | -3.027 | 0.002* | -0.438 | 0.345 |
|  | pine proximity | 0.411 | 0.234 | 1.758 | 0.079. | 0.217 |  |
|  | inorg. phosp. | 0.093 | 0.095 | 0.974 | 0.33 | 0.1 |  |
|  | soil pH | 0.187 | 0.114 | 1.641 | 0.101 | 0.203 |  |
|  | NH4+ | -0.397 | 0.094 | -4.202 | 0** | -0.424 |  |
|  | NO3- | 0.326 | 0.121 | 2.697 | 0.007* | 0.272 |  |
| inorg. phosp. | soil heating | 0.811 | 0.247 | 3.289 | 0.001** | 0.555 | 0.2 |
|  | pine proximity | -0.396 | 0.295 | -1.342 | 0.179 | -0.193 |  |
|  | fine fuel combustion | -0.487 | 0.143 | -3.399 | 0.001** | -0.486 |  |
| C:N ratio | pine proximity | 0.443 | 0.219 | 2.025 | 0.043* | 0.219 | 0.105 |
|  | fine fuel combustion | -0.213 | 0.106 | -2.007 | 0.045* | -0.215 |  |
| soil pH | soil heating | 1.05 | 0.197 | 5.319 | 0** | 0.719 | 0.507 |
|  | pine proximity | -1.167 | 0.244 | -4.786 | 0** | -0.57 |  |
| month 2 decomp. rate | litter fungi PCA 1 | -0.165 | 0.117 | -1.408 | 0.159 | -0.16 | 0.251 |
|  | plant comm. PCoA 1 | 0.341 | 0.131 | 2.611 | 0.009* | 0.402 |  |
|  | inorg. phosp. | -0.109 | 0.132 | -0.823 | 0.41 | -0.125 |  |
|  | C:N ratio | 0.198 | 0.112 | 1.77 | 0.077. | 0.226 |  |
|  | soil pH | -0.121 | 0.101 | -1.201 | 0.23 | -0.139 |  |
|  | NH4+ | 0.214 | 0.108 | 1.988 | 0.047* | 0.244 |  |
| month 4 decomp. rate | soil fungi PCA 1 | -0.212 | 0.099 | -2.144 | 0.032* | -0.21 | 0.341 |
|  | Soil fungi PCA 2 | 0.376 | 0.108 | 3.489 | 0** | 0.371 |  |
|  | Litter fungi PCA 1 | 0.679 | 0.163 | 4.158 | 0** | 0.587 |  |
|  | litter fungal diversity | -0.495 | 0.156 | -3.17 | 0.002** | -0.423 |  |
|  | plant comm. PCoA 1 | 0.174 | 0.095 | 1.828 | 0.068. | 0.183 |  |
|  | plant comm. PCoA 2 | 0.128 | 0.111 | 1.152 | 0.249 | 0.124 |  |
|  | C:N ratio | 0.331 | 0.102 | 3.256 | 0.001** | 0.337 |  |
| month 6 decomp. rate | soil fungi PCA 1 | 0.253 | 0.146 | 1.736 | 0.083. | 0.236 | 0.327 |
|  | soil fungi PCA 2 | 0.323 | 0.156 | 2.077 | 0.038* | 0.3 |  |
|  | soil fungal diversity | -0.264 | 0.151 | -1.747 | 0.081. | -0.239 |  |
|  | litter fungi PCA 2 | 0.11 | 0.114 | 0.962 | 0.336 | 0.095 |  |
|  | plant comm. PCoA 2 | 0.194 | 0.129 | 1.506 | 0.132 | 0.178 |  |
|  | C:N ratio | 0.512 | 0.118 | 4.329 | 0** | 0.491 |  |
|  | NH4+ | -0.154 | 0.106 | -1.454 | 0.146 | -0.148 |  |
|  | NO3- | 0.356 | 0.143 | 2.498 | 0.012* | 0.267 |  |
| month 8 decomp. rate | litter fungi PCA 1 | 0.208 | 0.119 | 1.745 | 0.081. | 0.192 | 0.22 |
|  | plant comm. PCoA 1 | 0.261 | 0.125 | 2.086 | 0.037* | 0.293 |  |
|  | plant comm. PCoA 2 | -0.152 | 0.105 | -1.45 | 0.147 | -0.159 |  |
|  | inorg. phosp. | -0.191 | 0.116 | -1.652 | 0.099. | -0.211 |  |
|  | C:N ratio | 0.272 | 0.106 | 2.554 | 0.011* | 0.296 |  |
|  | NO3- | 0.259 | 0.118 | 2.186 | 0.029* | 0.221 |  |
| Covariance structure | | Estimate | S.E. | z-value | p-value | Std. est. |  |
| soil fungi PCA 1 | soil fungal diversity | 0.397 | 0.092 | 4.292 | 0** | 0.629 |  |
| soil fungi PCA 2 | soil fungal diversity | 0.356 | 0.091 | 3.932 | 0** | 0.56 |  |
| litter fungi PCA 1 | litter fungal diversity | 0.503 | 0.1 | 5.044 | 0** | 0.807 |  |
| litter fungi PCA 2 | litter fungal diversity | -0.565 | 0.11 | -5.142 | 0** | -0.831 |  |
| litter fungi PCA 1 | litter fungi PCA 2 | -0.464 | 0.104 | -4.459 | 0** | -0.664 |  |
| plant comm. PCoA 1 | plant species richness | 0.297 | 0.074 | 3.986 | 0** | 0.572 |  |
| soil fungi PCA 1 | Soil fungi PCA 2 | 0.284 | 0.079 | 3.615 | 0** | 0.503 |  |
| inorg. phosp. | C:N ratio | 0.248 | 0.103 | 2.399 | 0.016* | 0.284 |  |
| C:N ratio | NH4+ | 0.347 | 0.119 | 2.912 | 0.004* | 0.359 |  |
| C:N ratio | NO3- | -0.142 | 0.084 | -1.699 | 0.089. | -0.189 |  |
| C:N ratio | soil pH | -0.106 | 0.077 | -1.382 | 0.167 | -0.155 |  |
| month 2 dec. rate | month 4 dec. rate | 0.082 | 0.077 | 1.063 | 0.288 | 0.133 |  |
| month 2 dec. rate | month 6 dec. rate | 0.017 | 0.082 | 0.204 | 0.838 | 0.025 |  |
| month 2 dec. rate | month 8 dec. rate | 0.044 | 0.078 | 0.571 | 0.568 | 0.071 |  |
| month 4 dec. rate | month 6 dec. rate | 0.306 | 0.094 | 3.242 | 0.001** | 0.439 |  |
| month 4 dec. rate | month 8 dec. rate | 0.378 | 0.094 | 4.007 | 0** | 0.573 |  |
| month 6 dec. rate | month 8 dec. rate | 0.198 | 0.091 | 2.177 | 0.029* | 0.281 |  |
| . = p<0.1; * = p<0.05; ** = p<0.0001 | | | | | | | |

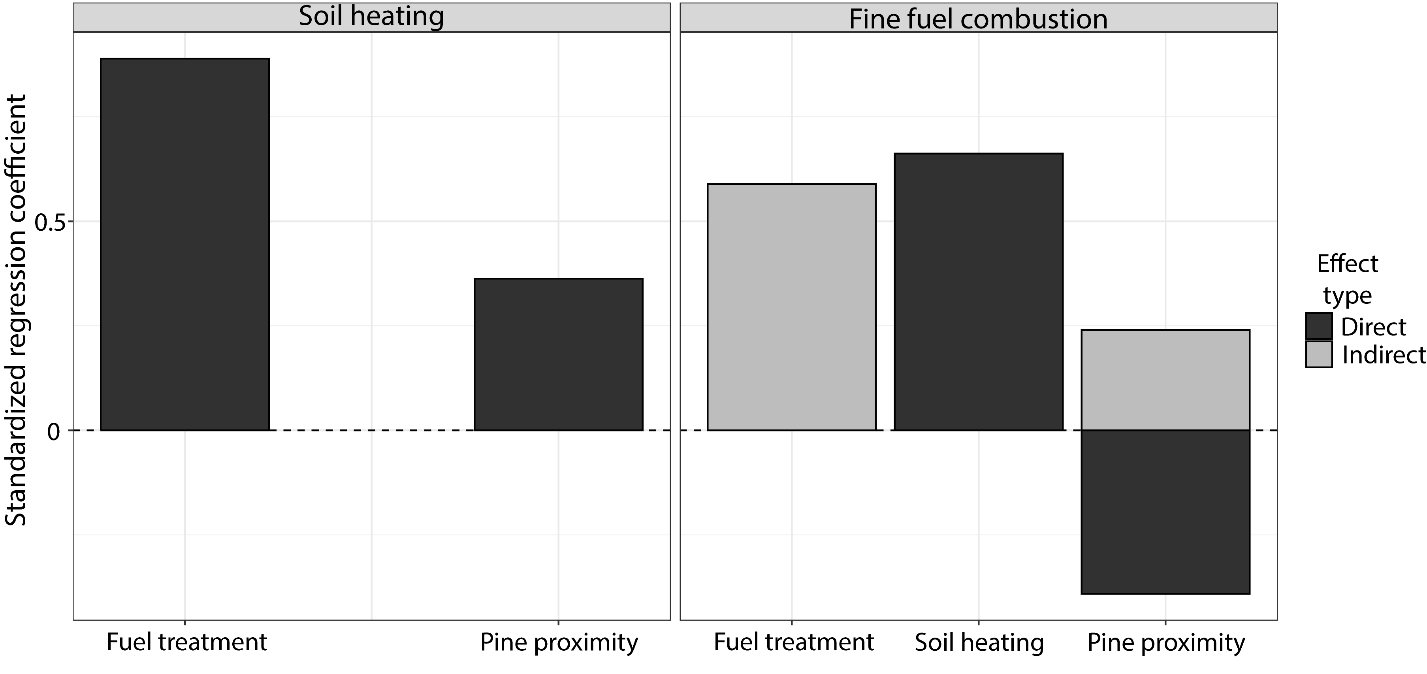

**Figure S31:** SEM variable effect sizes on soil heating and fine fuel combustion. Direct effects (black) represent paths linking the response and independent variable, while indirect effects (grey) represent paths between variables that are mediated by one or more variables.
